# Sex chromosome turnover and mitonuclear conflict drive reproductive isolation

**DOI:** 10.1101/2025.08.29.673102

**Authors:** Wynne V. Radcliffe, Mysia Dye, Daemin Kim, Pavitra Muralidhar, Rachel L. Moran

**Affiliations:** Department of Ecology & Evolution, University of Chicago, Chicago, IL, USA; Department of Biology, Texas A&M University, College Station, TX, USA; Department of Biology, Whitman College, Walla Walla, WA, USA

**Keywords:** reinforcement, speciation, sex chromosome turnover, mitonuclear conflict, hybrid incompatibility

## Abstract

Identifying the genetic basis of reproductive barriers is essential for understanding the origin and maintenance of biological diversity. While some hybrid incompatibilities evolve as incidental byproducts of divergence^1-3^, those involving sex chromosomes and mitochondrial-nuclear interactions may arise through predictable pathways shaped by genomic conflict^4-8^. Yet, the extent to which such interactions drive the evolution of reproductive barriers and speciation in natural populations remains unclear^9-11^. Here, we use whole-genome resequencing in North American fishes to show that two hybridizing species possess distinct, nonhomologous sex chromosomes. These chromosomes exhibit strong associations with sex, reduced introgression in natural hybrid zones, segregation distortion in backcrosses, and an enrichment of nuclear-encoded mitochondrial genes, indicative of sex-linked mitonuclear incompatibilities. We identify a third, distinct sex chromosome in another hybridizing species, indicating repeated sex chromosome turnover within the clade. Parental crosses and genomic analyses suggest that at least one of these transitions was driven by a recessive female-determining mutation, a rare empirical example of a theoretically predicted but seldom observed mechanism of sex chromosome evolution. Together, these results link genomic architecture to hybrid dysfunction and behavioral isolation, providing strong empirical support for long-standing predictions about the role of sex-linked and cytonuclear incompatibilities in speciation.

## INTRODUCTION

A major goal in evolutionary biology is to identify the genetic loci that contribute to reproductive isolation between species. Because reduced hybrid fitness can both initiate and reinforce reproductive barriers, uncovering its genetic basis is key to understanding the mechanisms and predictability of speciation. These barriers often originate during allopatric divergence and are revealed when lineages come into secondary contact. In such cases, hybrid dysfunction can drive the evolution of stronger prezygotic isolation (reinforcement), completing the speciation process and providing a direct link between intrinsic incompatibilities and the evolution of reproductive barriers^1^. While some hybrid incompatibilities evolve through stochastic divergence^2,3^, those involving the sex chromosomes and mitochondrial-nuclear (mitonuclear) interactions may evolve in predictable ways due to asymmetries in inheritance and selection^4,5^. Because sex chromosomes and mitochondrial DNA follow non-Mendelian inheritance patterns and often harbor rapidly evolving genes, they are predicted to be hotspots for hybrid incompatibilities^6,7^. Yet, few studies have directly tested these predictions with genomic data in natural systems with ongoing hybridization.

Sex chromosomes can exacerbate incompatibilities both directly and by compounding mitonuclear mismatches. The heterogametic sex (XY or ZW) often exhibits stronger hybrid dysfunction (Haldane’s rule)^8^, and mismatches between divergent sex chromosomes can generate epistatic incompatibilities that reduce hybrid fitness^9^. These effects may be further amplified when sex chromosomes accumulate mutations or structural variants under reduced recombination^10-12^. Moreover, nuclear genes that interact with mitochondria are frequently sex-linked and subject to different selective pressures in males and females^13^, raising the potential for compounded incompatibilities when sex chromosomes diverge in concert with mitonuclear coevolution^14-16^.

Teleost fishes exhibit remarkable diversity in sex determination systems, making them especially valuable for investigating the evolutionary consequences of sex chromosome turnover. Even within a genus, sex chromosomes can vary in structure, gene content, and identity^17,18^. Environmental influences, such as temperature, can also affect sex determination^19-22^. This variability complicates predictions about hybrid outcomes but offers a unique opportunity to examine the interplay between sex chromosome evolution and reproductive isolation.

Darters (Percidae: Etheostominae) are a diverse radiation of North American stream fishes that have emerged as a powerful model for studying the evolutionary mechanisms that drive species diversification^23,24^. Divergence is typically initiated in allopatry in darters, but many species hybridize in sympatry after secondary contact^25,26^. Reinforcement has been inferred in several lineages^23,27,28^, most notably between two members of the subgenus *Oligocephalus*, orangethroat darters (*Etheostoma spectabile*) and rainbow darters (*Etheostoma caeruleum*) which readily hybridize in sympatry. These taxa exhibit stronger behavioral isolation in sympatry than in allopatry^23,27^ and produce low-fitness hybrids characterized by male-skewed sex ratios and reduced backcross viability^24, 28^. These patterns suggest that intrinsic genetic incompatibilities may contribute to reproductive isolation and that such incompatibilities could have promoted the evolution of enhanced prezygotic isolation in sympatry.

Here, we investigate the genomic basis of sex determination and its contribution to reproductive isolation in darters using whole-genome resequencing and population genomic analyses. We first focus on *E. spectabile* and *E. caeruleum* and show that these hybridizing species possess distinct, nonhomologous sex chromosomes, consistent with independent sex chromosome turnovers prior to secondary contact. Both species exhibit male heterogamety, and we identify candidate sex-determining genes in regions of elevated X-Y divergence. These chromosomes show reduced introgression, segregation distortion in hybrids, and an enrichment of nuclear-encoded mitochondrial genes in sex-associated regions, patterns consistent with sex-linked mitonuclear incompatibilities. Biparental crosses also suggest that this turnover was driven by a recessive female-determining mutation, representing a rare empirical example of a theoretically predicted but seldom documented pathway of sex chromosome evolution^29,30^. To place these findings in a broader phylogenetic context, we extended our analyses to additional hybridizing darter species within the *Oligocephalus* subgenus. This comparative framework reveals both conservation of sex chromosome identity within some lineages and additional, independent turnovers in others, demonstrating that sex chromosome evolution is a recurrent and lineage-specific phenomenon in this clade. Together, our findings demonstrate how dynamic sex chromosome evolution and cytonuclear conflict intersect to generate hybrid incompatibilities, offering new insight into the predictability of reproductive barriers and the genomic mechanisms that facilitate the maintenance of species boundaries in the presence of hybridization.

## RESULTS

### Etheostoma caeruleum genome assembly

We previously sequenced and assembled a chromosome-level reference genome for *E. spectabile* (GCF_008692095.1)^24^. To prevent reference bias in our population genetic analyses, we generated a *de novo* reference genome for *E. caeruleum* (see Supplementary Methods, Supplementary Table 1). The new assembly showed high completeness as indicated by BUSCO scores (C: 92.3% [S:91.6%, D:0.7%], F:3.9%, M:3.8%) and repeat content (40.45% GC content, 33.02% bases masked) comparable to other published *Etheostoma* genomes, including *E. spectabile*, *E. cragini*, and *E. perlongum.* Our *E. caeruleum* reference genome is largely syntenic with the *E. spectabile* reference genome, with high collinearity between the two (Extended Data Figure 1).

### Sex-linked loci identified through genome-wide association

We conducted genome-wide association studies (GWAS) using GEMMA (v0.98.5) to detect loci associated with sex in both *E. spectabile* and *E. caeruleum*. To evaluate the broader evolutionary dynamics of sex chromosome identity in this clade, we extended our GWAS analyses to two additional hybridizing species within *Oligocephalus*: *E. pulchellum* (a member of the orangethroat darter complex, *Ceasia*) and *E. radiosum* (Figure 1a). Our analysis identified a strong sex association on chromosome 9 in *E. spectabile* (Figure 1b). Similarly, a significant peak was observed on chromosome 23 in *E. caeruleum* (Figure 1b). When analyzed separately, allopatric populations exhibited a stronger association signal on the sex chromosome compared to sympatric populations for the *E. caeruleum* (Extended Data Figs. 2-3). Analysis of intersex F_ST_ revealed consistent patterns (Extended Data Figure 2). This suggests that divergence in the sex-determining region may have predated secondary contact. In *E. pulchellum*, our analyses revealed support for chromosome 9 as the sex chromosome (Extended Data Fig. 4), indicating potential sex chromosome conservation within *Ceasia*. However, we identified a unique sex-associated region on chromosome 22 in *E. radiosum* (Figure 1b). This indicates repeated, independent sex chromosome turnover events have occurred within *Oligocephalus* (Figure 1c).

**Figure 1.**
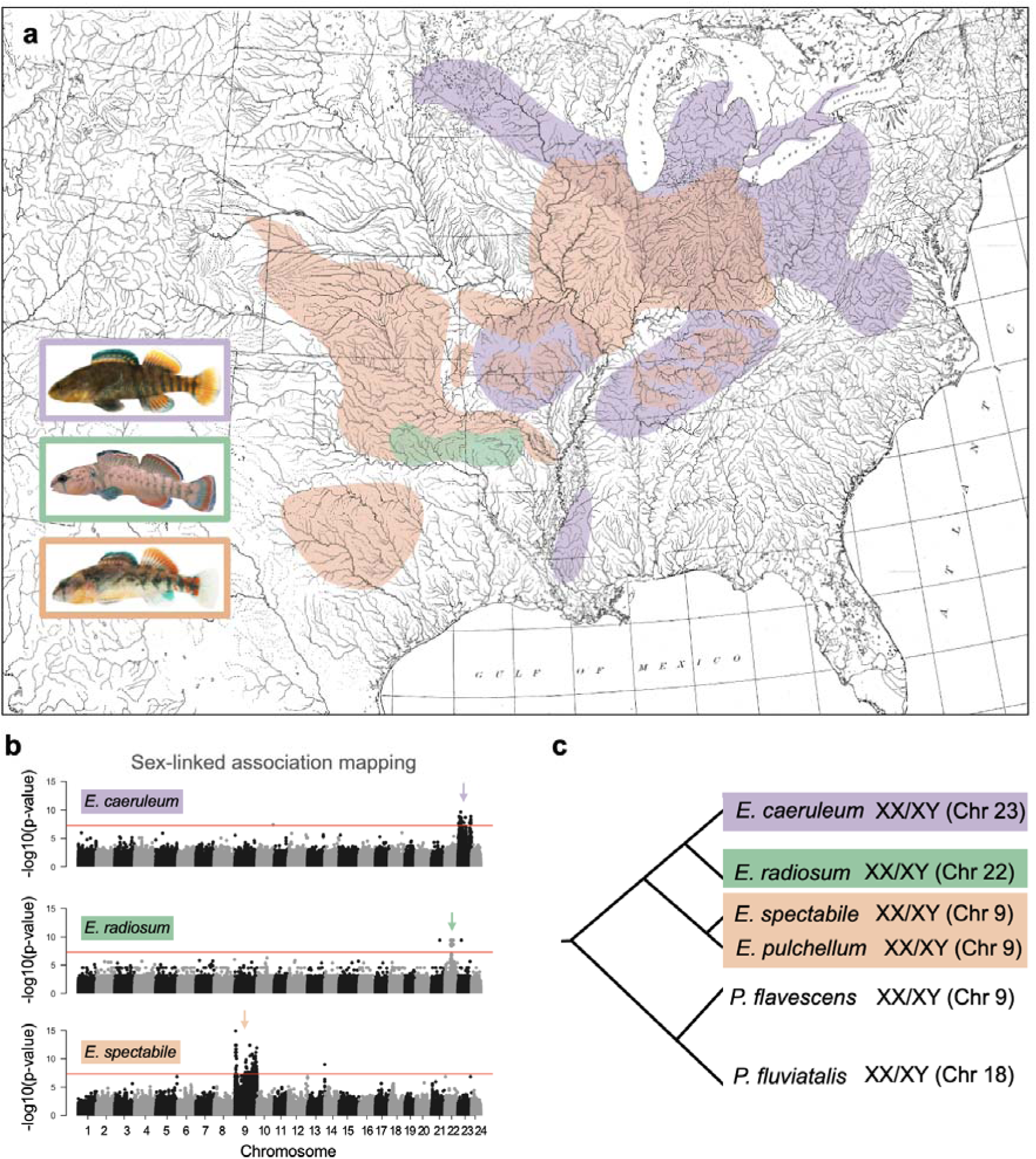
Repeated sex chromosomes turnover between hybridizing species. (a) Range maps for *E. caeruleum* (purple), the *E. radiosum* species complex (green), and the *E. spectabile* species complex (orange; includes *E. pulchellum*). (b) Results of genome-wide association study (GWAS) for sex in *E. caeruleum*, *E. radiosum*, and *E. spectabile* showing sex chromosome turnover has occurred repeatedly in darter lineages that now co-occur and hybridize in sympatry, suggesting a potential role in reinforcing species boundaries. GWAS revealed strongest differentiation between sexes on chromosome 23 in *E. caeruleum*, chromosome 22 in *E. radiosum*, and chromosome 9 in *E. spectabil*e and *E. pulchellum* (see Extended Data Figure 4). (c) Phylogeny showing sex chromosome turnover in darters and non-darter percids. Chromosome 9 is the putative ancestral sex chromosome, shared by *Perca flavescens* and multiple members of the orangethroat darter complex (e.g., *E. spectabile*, *E. pulchellum*).

### Sex-linked differences in heterozygosity and SNP density

To characterize sex-linked patterns of variation, we compared genome-wide heterozygosity and SNP density between males and females. In species with male heterogamety (XY), males are expected to show elevated heterozygosity on the sex chromosomes relative to autosomes. Consistent with this expectation, *E. spectabile* showed strong male-biased heterozygosity restricted to chromosome 9 (Wilcoxon rank-sum, *p* < 1 × 10^-43^; Cliff’s δ = 0.60, large effect; Supplementary Table 2, Fig. 2a), and *E. caeruleum* showed strong male-biased heterozygosity restricted to chromosome 23 (*p* < 1 × 10^-35^; Cliff’s δ = 0.72, large effect; Supplementary Table 2, Fig. 2b), each consistent with an XY system. In *E. radiosum*, chromosome 22 displayed more modest but still significant male-biased heterozygosity (*p* < 1 × 10^-14^; Cliff’s δ = 0.44, large effect; Supplementary Table 2, Fig. 2c), suggesting that recombination suppression and divergence between the X and Y are more limited or recent in this lineage.

**Figure 2.**
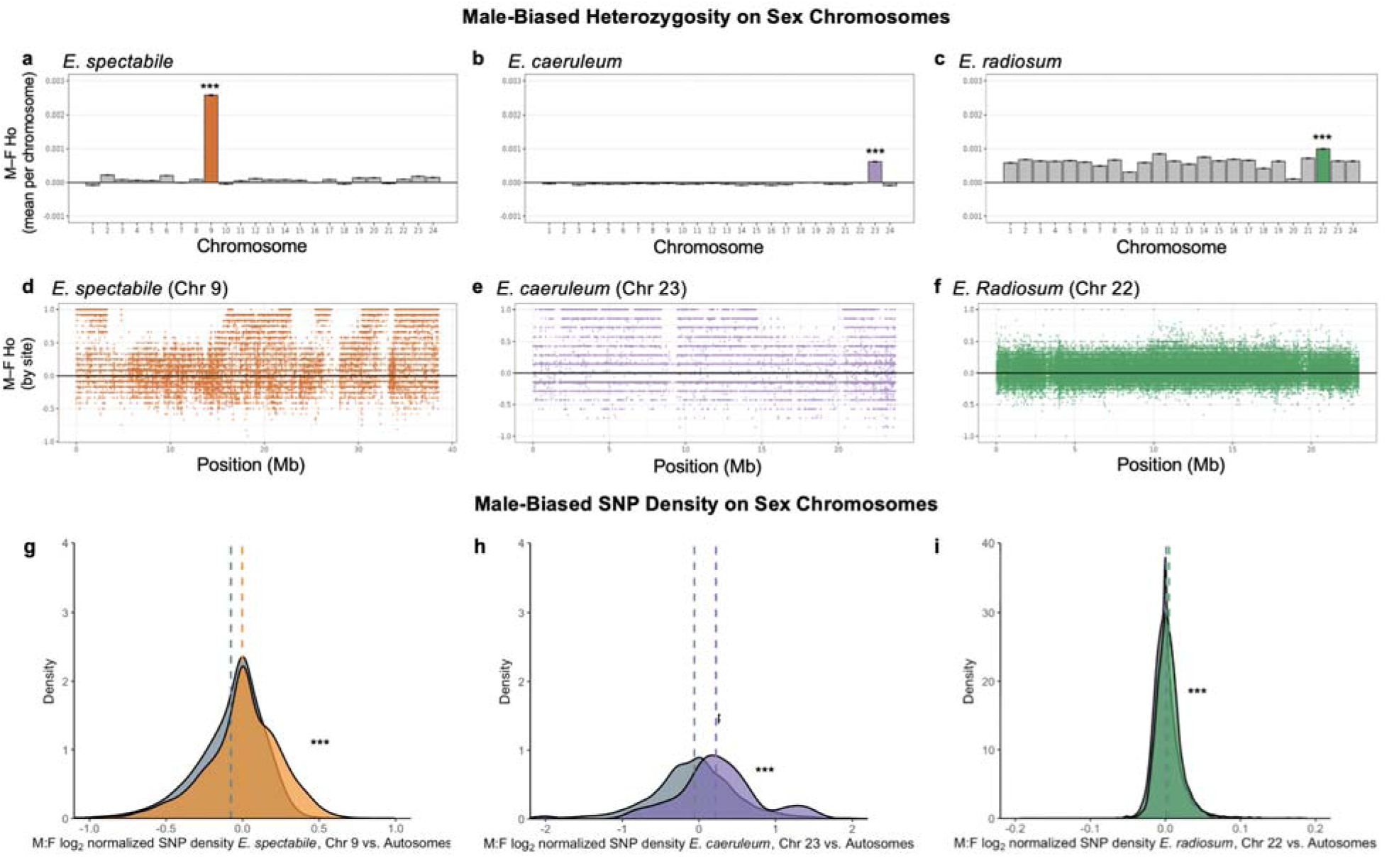
Male-biased heterozygosity and SNP density reveal XY heterogamety across darters with independent sex chromosome turnovers. Panels show patterns in *E. spectabile* (left), *E. caeruleum* (middle), and *E. radiosum* (right). Top row (a–c): mean ± SE of male–female heterozygosity differences (M–F Ho) across each of the 24 chromosomes. Sex-linked chromosomes showed significantly elevated male-biased heterozygosity (*E. spectabile*: Wilcoxon rank-sum, *p* < 1 × 10^-43^, Cliff’s δ = 0.60; *E. caeruleum*: *p* < 1 × 10^-35^, δ = 0.72; *E. radiosum*: *p* < 1 × 10^-14^, δ = 0.44). Middle row (d–f): site-level M–F heterozygosity differences along the identified sex chromosome in each species. Bottom row (g–i): distributions of male:female log□-normalized SNP density for autosomes (grey) versus sex chromosomes (colored). Again, sex-linked chromosomes showed elevated male-biased SNP density (*E. spectabile*: *p* < 1 × 10^-15^, δ = 0.12; *E. caeruleum*: *p* = 3.3 × 10^-6^, δ = 0.31; *E. radiosum*: *p* < 1 × 10^- 15^, δ = 0.12). Together these results demonstrate parallel evolution of XY systems across darters, with conserved male heterogamety but species-specific differences in the extent and genomic footprint of X–Y divergence (ancestral chromosome 9 in *E. spectabile*; neo-sex chromosome 23 in *E. caeruleum*; neo-sex chromosome 22 in *E. radiosum*). Asterisks indicate significance thresholds in panel a–c and g-i (*** = *p* < 1 × 10^-5^).

Analyses of SNP density corroborated these patterns across all three species (Fig. 2g–i). In *E. spectabile*, autosomal SNP densities were tightly distributed around zero, while chromosome 9 showed a slight but significant male bias, consistent with an established XY system (Wilcoxon rank-sum, *p* < 1 × 10^-15^; Cliff’s δ = 0.12, small effect; Supplementary Table 2, Fig. 2g). In *E. caeruleum*, the distribution of SNP density was broader overall, but chromosome 23 showed pronounced and widespread male bias, consistent with extensive differentiation of the neo-Y (Wilcoxon rank-sum, *p* = 3.3 × 10^-6^; Cliff’s δ = 0.31, medium effect; Supplementary Table 2, Fig. 2h). By contrast, *E. radiosum* exhibited a narrow distribution of autosomal SNP density and a more modest but still significant male bias on chromosome 22 (Wilcoxon rank-sum, *p* < 1 × 10^-15^; Cliff’s δ = 0.12, small effect; Supplementary Table 2, Fig. 2i), again consistent with more localized or younger sex chromosome divergence. Together, these results demonstrate parallel evolution of XY systems across darters, with species differing in the extent and genomic footprint of X–Y divergence.

These results, together with prior karyotype evidence of homomorphic sex chromosomes in darters^32^, point to chromosome 9 as the ancestral sex chromosome and to chromosomes 23 (*E. caeruleum*) and 22 (*E. radiosum*) as independently derived neo-sex chromosomes. Genome alignment further reveals that chromosomes 9 and 23 are largely syntenic between the *E. spectabile* and *E. caeruleum* reference genomes (Extended Data Fig. 1), supporting the hypothesis that chromosome 23 evolved as a neo-sex chromosome *de novo* in *E. caeruleum*, rather than by translocation of the ancestral sex-determining region.

### Recessive sex chromosome transition explains male-bias in F1 crosses

We find that a heterologous transition, induced by a recessive female-determining mutation, would account for the maintenance of male heterogamety in *E. spectabile* and *E. caeruleum*, while also explaining the previously observed male-biased sex ratios in F1 hybrid crosses^28^ (Extended Data Table 2, Fig. 3). Sex chromosome transitions involve a change from a recurrent pair of sexual genotypes – that is, a pair of sexual genotypes that create only themselves when mated to each other (e.g. XX and XY individuals) – to another stable recurrent pair^29,30^, This transition occurs via the creation of intermediate sexual genotypes, which create an evolutionary ‘pathway’ connecting these recurrent pairs, and thereby facilitating a transition from the ancestral recurrent sexual genotypes to the new recurrent sexual genotypes^33^. Evolution along the pathway connecting these recurrent pairs of sexual genotypes can be facilitated or hindered by drift^33,34^, or by selection for or against new sexual genotypes^35-37^. Because drift-induced or selective forces will tend to ‘push’ a population to one pair of recurrent sexual genotypes or the other, intermediate sexual genotypes are generally assumed to be transient, although the long-term within-population persistence of intermediate sexual genotypes has been noted in other fish species^38,39^.

**Figure 3.**
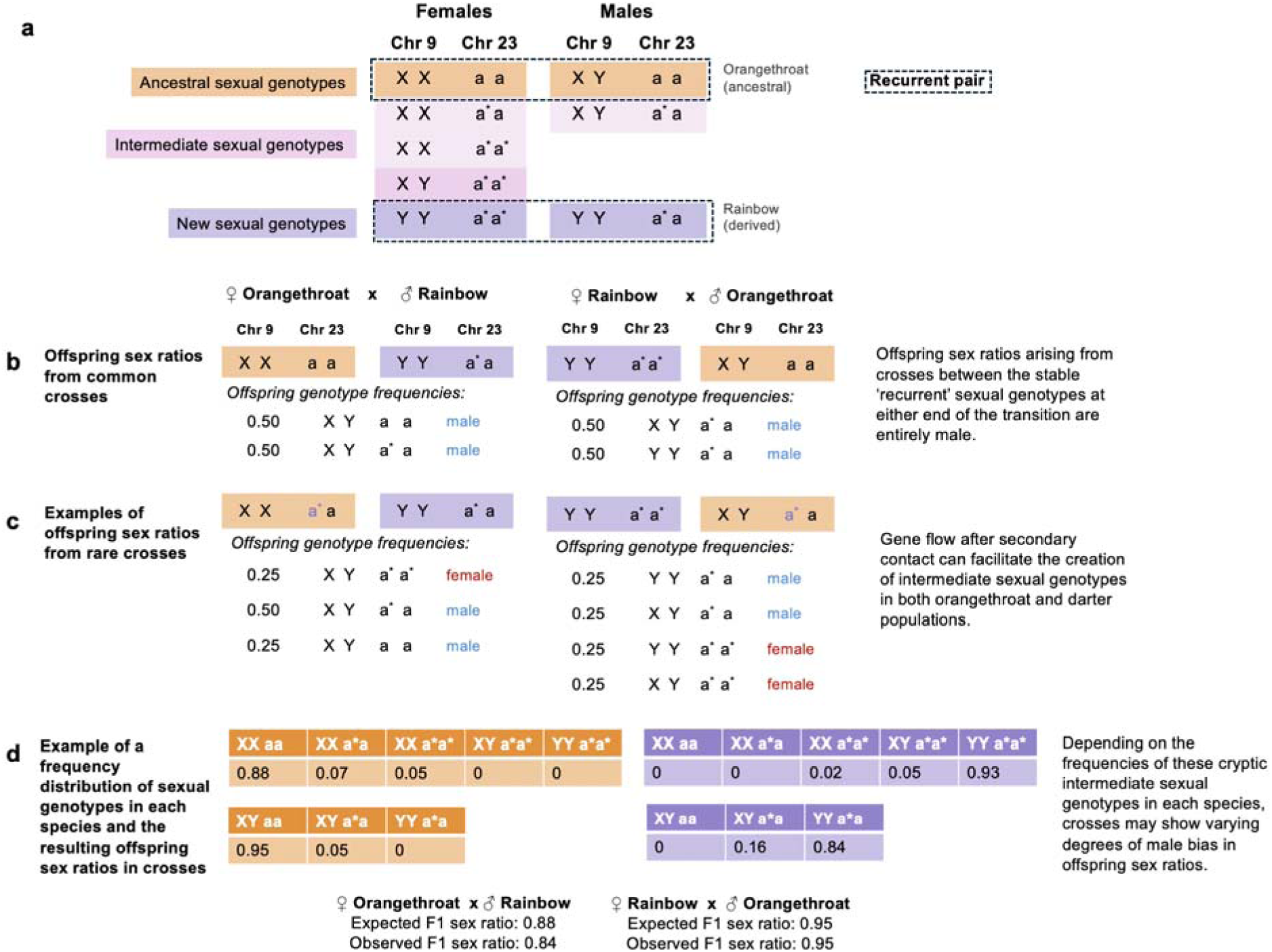
Sex chromosome turnover in darters via a ‘recessive’ heterologous transition. (a) The appearance of a recessive new female-determining mutation, ‘a*’, on autosome 23 in allopatry between *E. spectabile* (orangethroat darters) and *E. caeruleum* (rainbow darters), can lead to a sex chromosome turnover in *E. caeruleum*. This transition will establish chromosome pair 23 as the sex chromosomes in *E. caeruleum* (with the ‘a*’ haplotype acting as a neo X and ‘a’ acting as a neo-Y), and maintains male heterogamety. (b) Biparental crosses between the established sexual genotypes (the ‘recurrent pair’) in *E. spectabile* and *E. caeruleum* and will generate completely male-biased sex ratios in both directions of the cross. (c) Ongoing gene flow between *E. spectabile* and *E. caeruleum* populations will facilitate matings between individuals using either the ancestral or the derived sex chromosome system. This hybridization will recreate the intermediate sexual genotypes involved in the transition between the two sex chromosome systems. The offspring sex ratios of matings involving these intermediate sexual genotypes varies; two examples of such crosses are shown here for illustration. Full details of the predicted offspring sex ratios between all sexual genotypes are available in Extended Data Table 2. (d) While the full distribution of the frequencies of intermediate sexual genotypes in *E. spectabile* and *E. caeruleum* is unknown, different frequency distributions of the cryptic intermediate sexual genotypes in each species (and/or in different populations within each species) could generate a range of male-biased sex ratios consistent with the sex ratios observed in our biparental crosses. Here we show an example of the potential frequency distributions of sexual genotypes in each species, along with the resulting offspring sex ratios in biparental crosses between species given those distributions.

In the *E. spectabile* and *E. caeruleum* system, *E. caeruleum* appear to have undergone a recessive transition to a new male heterogametic system on chromosome 23, replacing the ancestral system on chromosome 9 (Fig. 3). If both species contained only the recurrent sexual genotypes found at either end of this transition (Fig. 3a), we would predict completely male-biased sex ratios in both directions of crosses between those species (Fig. 3b). We suggest, however, that ongoing gene flow between *E. spectabile* and *E. caeruleum* is facilitating the persistence of intermediate sexual genotypes in both species. Essentially, the ongoing hybridization between *E. spectabile* and *E. caeruleum* acts as a force constantly pulling these species away from the stable recurrent genotypes and creating – via matings with individuals from the sister species with a diverged sex chromosome system – a constantly refreshing stock of intermediate sexual genotypes.

While the sex ratios from crosses between *E. spectabile* and *E. caeruleum* are still predicted to be highly male-biased, the presence of these intermediate sexual genotypes will modulate the degree of male bias expected in the offspring of those crosses (Fig. 3c). The exact offspring sex ratios in each direction of the cross will depend on the frequencies of the intermediate genotypes in each species, which in turn will be sensitive to the level of gene flow and selection on introgressed ancestry across *E. spectabile* and *E. caeruleum* populations. However, in general, the resulting sex ratios will be highly, but not completely, male-biased in both directions of the cross, and can generate offspring sex ratios broadly consistent with the observed data from our crosses (Fig. 3d). Indeed, the biased sex ratios arising from matings between individuals from different species could even act as an additional force selecting against hybrids^24, 28^.

### Candidate genes involved in sex determination and mitochondrial function

We next looked for candidate sex determining genes on the sex chromosomes. We identified 219 genes within 50 kb of top SNPs from the GWAS for sex on chromosome 9 in orangethroat darters (see Supplementary Table 3) and 206 genes within 50 kb of top SNPs from the GWAS for sex on chromosome 23 in *E. caeruleum* (see Supplementary Table 3). A subset of these genes overlapped with known sex determining or mitochondrial functions.

In *E. spectabile,* sex-associated SNPs on chromosome 9 cluster across six regions of the zebrafish genome (chromosomes 6, 8, and 22), suggesting complex synteny relationships and potential reorganization of sex-associated loci across lineages (Extended Data Fig. 5). Candidate genes on chromosome 9 were enriched for ontologies related to germline stem cell maintenance, sperm chromatin condensation, meiotic division, and luteinizing hormone response. These genes included known sex determination loci in teleosts: *amh*, *tgfbr3*, *dmrta2*, *dmrt2b*, *sox14*, *sox2*, *vtg3*, and *brdt*^40,41^(Figure 4a).

**Figure 4.**
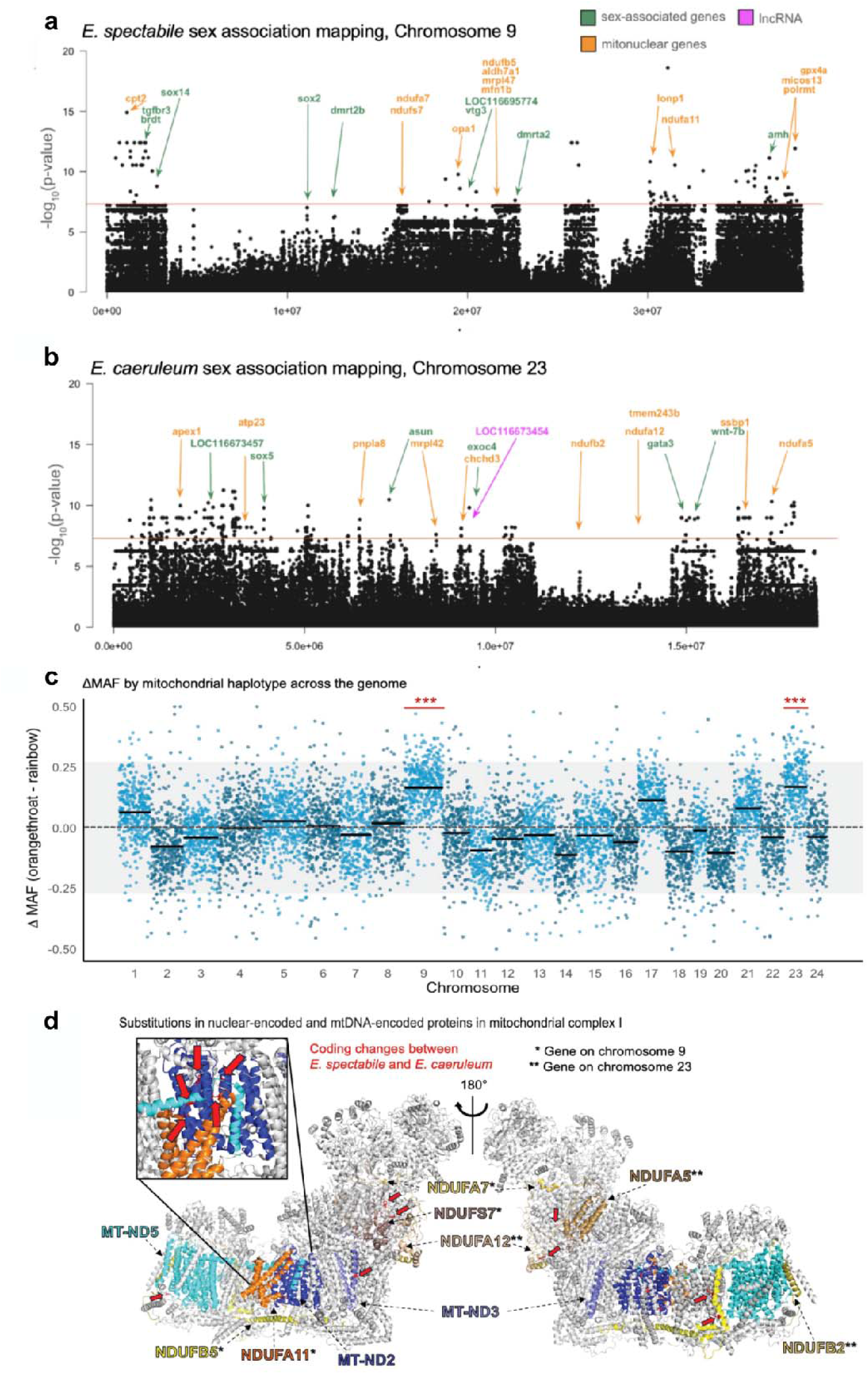
Sex chromosomes are hotspots for mitonuclear conflict. (a–b) Mitonuclear genes are shown in green, sex-associated genes in orange, and a long non-coding RNA (lncRNA) located within an exon is highlighted in pink. (a) In *E. spectabile*, chromosome 9 harbors ten mitonuclear genes and multiple sex-associated genes located within 50 kb of highly significant GWAS SNPs. LOC116695774 is predicted to be a vitellogenin-like pseudogene. (b) In *E. caeruleum*, chromosome 23 contains nine mitonuclear genes and four sex-associated genes within 50 kb of GRAS significant SNPs. LOC116673457 encodes a protein similar to fish-egg lectin, and LOC116673454 is a putative lncRNA predicted to target *amhr2* and *gnrhr.* (c) Changes in minor allele frequencies (ΔMAF) between backcrosses to femal*e E. spectabile* (*E. spectabile* mitochondrial genome) and backcrosses to female *E. caeruleum* (*E. caeruleum* mitochondrial genome) are elevated on sex chromosomes compared to the genome-wide expectation (*** = P<0001, Wilcoxon rank sum tests). The dashed dark gray line indicates the null expectation (ΔMAF = 0) and the shaded light grey band represents the 95% confidence interval. (d) Structural model of mitochondrial Complex I (PDB: 5XTD, visualized in PyMOL) highlights nuclear-encoded subunits located on chromosome 9 (NDUFS7, NDUFA7, NDUFA11, NDUFB5; *), chromosome 23 (NDUFA5, NDUFA12, NDUFB2; **), and their mitochondrial-encoded interacting proteins (MT-ND2, MT-ND3, MT-ND5). Residues with amino acid substitutions derived in *E*. *spectabile* and/or *E. caeruleum* are shown in red, based on comparative alignments across darter species^44^. Substitutions in NDUFS7, NDUFA11, NDUFB5, MT-ND2, MT-ND3, and MT-ND5 have been previously identified as targets of positive selection in darters and other percids^43^. Inset shows a detailed view of coding changes in three interacting subunits, NDUFA11, MT-ND2, and MT-ND5.

GO analysis also revealed an enrichment of genes associated with mitochondrial function in sex-associated regions on chromosome 9, including mitochondrial inner membrane fusion, mitochondrial transcription, mitochondrial DNA metabolic process, and response to oxidative stress (see Supplementary Table 4). We identified nine nuclear-encoded mitochondrial genes located in sex-associated regions on chromosome 9, including *ndufs7*, *ndufa7*, *ndufb5*, and *ndufa11*, which contribute to complex I of the oxidative phosphorylation (OXPHOS) pathway (Figure 3d). Additional genes implemented in mitochondrial DNA replication, protein import, fusion, and antioxidant activity, functions critical to mitochondrial integrity and signaling were also present in sex-associated regions on chromosome 9 (Figure 3d)^94^.

In *E. caeruleum*, top sex-associated SNPs on chromosome 23 were associated with genes enriched for vitellogenesis (Supplementary Table 4) and mapped to the right arm of zebrafish chromosome 4 (Chr4R), a known sex-associated region in *Danio rerio* (Extended Data Fig. 5). One of the most highly sex-associated SNPs overlapped a long non-coding RNA (lncRNA). A blastN search of the lncRNA coding region revealed high-confidence hits (≥85% identity, E-value <1e-5) to genes such as *CLDN34*, *amhr2*, *gnrhr1*, *gnrhr2*, and *rsrc1*. The homology to genes encoding anti-mullerian hormone receptor (*amhr2*) and gonadotropin releasing hormone receptors (*gnrhr1, gnrhr2*) suggests that this lncRNA may play a regulatory role in sex determination pathways.

Genes with mitochondrial-related functions were also enriched within highly sex-associated regions on chromosome 23, again including genes associated with complex I of the oxidative phosphorylation (OXPHOS) pathway (*ndufa5*, *ndufb2*, and *ndufa12*) (Supplementary Table 4, Figure 4b). Other genes in highly sex-associated regions of chromosome 23 were enriched for ontologies related to respiratory chain assembly, NAD metabolism, phosphate transport, and tRNA modification (Supplementary Table 4). Additional enriched GO categories included dopaminergic neuron differentiation, glutamatergic transmission, and visual learning, potentially linking sex determination to broader neural or behavioral traits.

In *E. radiosum*, we identified a novel sex-associated region on chromosome 22 containing candidate genes involved in neuroendocrine signaling and mitochondrial function, including *shh*, *oprk1*, *ptprn2*, *adcyap1*, and *ndufb4*, a nuclear-encoded mitochondrial complex I subunit (see Supplementary Note). The presence of *ndufb4* supports a recurring pattern across darters in which independently evolved sex chromosomes harbor genes mediating hormonal and cytonuclear interactions, potentially reflecting selection for restored mitonuclear compatibility in hybridizing lineages.

### Ancestry inference reveals resistance to introgression in sex-linked regions

To test whether sex chromosomes are resistant to introgression, we used two complementary approaches. First, TWISST analyses revealed that genomic regions with strong sex associations also had higher support for the species tree topology (Extended Data Fig. 6), indicating reduced phylogenetic discordance and gene flow (*E. spectabile* chromosome 9: r = 0.13, p < 2.2e^−16^; *E. caeruleum* chromosome 23: r = 0.06, p < 2.2e^−16^; Extended Data Fig. 6). Second, *fdM* statistics were significantly lower on sex chromosomes than autosomes (Wilcoxon rank-sum test: W = 2, p = 0.029), consistent with reduced allele sharing (Supplementary Table 5). Backcrosses further support this pattern: chromosomes 9 and 23 showed elevated segregation distortion, with chromosome 23 displaying complete absence of recombinant haplotypes in backcrosses to *E. caeruleum* females (p < 0.001)^24^. These results suggest strong selection against recombination on chromosome 23, potentially due to lethal mitonuclear incompatibilities involving nuclear-encoded mitochondrial genes linked to the Y chromosome.

### Mitonuclear associations map to the sex chromosomes

To identify nuclear loci whose allele frequencies depend on mitochondrial background, we tested for deviations from the expected minor allele frequency (MAF = 0.25) across all SNPs in *E. spectabile* x *E. caeruleum* backcross hybrids. Genome-wide chi-square tests identified 558 out of 8,177 SNPs with significant associations between mitochondrial haplotype and nuclear genotype. Three chromosomes harbored more mitonuclear associations than expected by chance: chromosome 9 (odds ratio = 2.88, FDR-adjusted p = 2.81 × 10^-9^), chromosome 18 (odds ratio = 2.02, FDR-adjusted p = 0.004), and chromosome 23 (odds ratio = 3.68, FDR-adjusted p = 1.02 × 10^-9^). Notably, the sex chromosomes, 9 and 23, harbored the greatest number of significant SNPs (Supplementary Table 6), suggesting these chromosomes are disproportionately involved in mitonuclear interactions in hybrids.

We further quantified allele frequency shifts between mitonuclear backgrounds by calculating ΔMAF for each SNP (ΔMAF = MAF in hybrids with *E. spectabile* mitochondria − MAF in hybrids with *E. caeruleum* mitochondria). Under a null model of no mitonuclear effects, ΔMAF values should be centered around zero. Instead, mean ΔMAF was significantly elevated on chromosomes 9 (ΔMAF = 0.166) and 23 (ΔMAF = 0.169) (Figure 4c). Permutation tests confirmed that these values were greater than expected by chance (both p < 1×10^-4^). Wilcoxon tests further indicated that the distributions of ΔMAF values on chromosome 9 (p = 5.9×10^-154^) and chromosome 23 (p = 8.7×10^-93^) were significantly skewed compared to the rest of the genome. The positive value of ΔMAF indicates that *E. caeruleum* alleles are present at elevated frequencies on chromosomes 9 and 23 in backcrosses to *E. spectabile* (relative to backcrosses to *E. caeruleum*). Notably, mitonuclear genes are evenly distributed across all chromosomes (Supplementary Table 6). This suggests that although mitonuclear genes are genome-wide, those located on sex chromosomes contribute disproportionately to hybrid incompatibilities.

Genes within 50 kb of SNPs significantly associated with mitochondrial haplotype were enriched for functions related to male sex differentiation, sperm capacitation, mitochondrial membrane organization, and mitochondrial DNA replication and metabolism. These GO terms reinforce the functional intersection of sex-specific and mitochondrial processes in regions implicated in incompatibilities.

### Functional implications of mitonuclear substitutions

Previous work has identified derived substitutions in mitochondrial-encoded genes (e.g., *mt-nd2, mt-nd3, mt-nd5*) and evidence of positive selection in nuclear OXPHOS genes in darters^43,44^. Here, we identified derived coding changes in both *E. spectabile* and *E. caeruleum* in nuclear-encoded complex I genes located on the sex chromosomes (*ndufa7*, *ndufa11*, *ndufa12*, *ndufb5*, *ndufs7*). *In silico* functional predictions with SIFT^95^ suggest that two variants, P45S in NDUFS7 (chromosome 9) and Q116R in NDUFA12 (chromosome 23), both derived in *E. spectabile*, are likely deleterious (Extended Data Table 3). These residues are otherwise conserved across darter and perch species^44^ (Extended Data Fig. 7), supporting the hypothesis that these changes may contribute to mitonuclear incompatibilities in hybrids.

## DISCUSSION

Our study reveals that sex chromosome turnover and mitonuclear incompatibilities jointly shape the genomic landscape of reproductive isolation in darters. We identify three nonhomologous sex chromosomes in hybridizing species within *Oligocephalus*, indicating multiple independent sex chromosome transitions. Our findings provide compelling evidence for a recessive female-determining mutation underlying one of the turnovers, offering a rare empirical example of a theoretically anticipated but rarely observed mode of sex determination evolution. These divergent sex chromosomes are associated with segregation distortion and enrichment of nuclear-encoded mitochondrial genes, especially from the OXPHOS pathway. Our findings suggest that mismatches between sex-determining systems and disrupted mitonuclear coevolution contribute to hybrid dysfunction and reinforce species boundaries. Together, these results highlight sex chromosomes as dynamic hotspots of evolutionary conflict and speciation.

### Sex chromosome divergence and hybrid incompatibility

Our findings suggest that incompatibilities between divergent sex-determining systems contribute substantially to postzygotic isolation in darters. While *E. spectabile* and *E. caeruleum* both show male-heterogamety (XY), each species utilizes a different pair of chromosomes as the sex chromosomes: *E. spectabile* use chromosome 9, the ancestral sex chromosome system in this clade, while *E. caeruleum* use chromosome 23. Chromosome 9 likely represents the ancestral sex chromosome in this clade, because it is also the sex chromosome in yellow perch (*Perca flavescens*)^42^, an earlier diverging lineage in Percidae^43^. Notably, while the yellow perch Y-specific gene *amhr2by* is absent in darters, the chromosomal substrate appears conserved. The novel chromosome 23 sex-determining region in *E. caeruleum* includes *asun*, a gene involved in spermatogenesis^45^, which may have facilitated the emergence of a new sex-determining mutation in this region. Note that, although this gene is involved in spermatogenesis, a loss-of-function mutation at this gene could potentially still act as a recessive female-determining mutation^45-47^.

We have presented evidence, via parental crosses and analyses of sex chromosome transitions, that there was likely a heterologous transition in sex determination between these species, and that this transition involved the invasion of a ‘recessive’ female sex-determining mutation in *E. caeruleum*. While transitions induced by both dominant and recessive sex-determining mutations have been well-characterized theoretically^33,34^, transitions in sex determination induced by dominant mutations are predicted to be more common^35, 36^ and have been better characterized empirically^48,49^. The genomic characterization of a recessive sex chromosome turnover is therefore a key contribution to the growing literature on the evolutionary lability of sex-determining mechanisms across vertebrates^50-53^.

We have further postulated that there exist cryptic intermediate sexual genotypes in each species, based on the highly, but not completely, male-biased sex ratios observed in biparental crosses^28^. The persistence of these intermediate genotypes is likely facilitated by ongoing gene flow between the two parental species^28^, as matings between individuals from populations with different sex chromosome systems will regenerate the intermediate genotypes involved in the transition between these sex chromosome systems. Further investigation will be required to establish the prevalence of these cryptic intermediate sexual genotypes in *E. spectabile* and *E. caeruleum*, and the fitness consequences of this intraspecific variation in genotypic sex determination.

### Turnover and reinforcement as a driver of speciation

Sex chromosomes evolve rapidly across animals, and their divergence is often implicated in reproductive isolation and speciation. While most insights come from systems with highly differentiated sex chromosomes (e.g., mammals, birds), teleost fishes exhibit exceptional diversity in sex-determining systems. We observed that *E. spectabile/E. puchellum*, *E. caeruleum, and E. radiosum* each uses a different sex chromosome (Fig. 1), and patterns of sex-association in allopatry versus sympatry suggest that these turnovers occurred prior to secondary contact (Extended Data Figs. 2-3). In *E. caeruleum*, the signal of sex linkage on chromosome 23 is substantially reduced in sympatric populations, possibly due to gene flow eroding linkage between the sex-determining region and male-biased markers. These observations suggest incompatibilities that evolved in allopatry may act as postzygotic barriers upon secondary contact, promoting the evolution of prezygotic isolation via reinforcement^28^.

Cross-taxon surveys underscore that rapid, lineage-specific sex-chromosome turnover is not unique to darters. Recurrent transitions have been documented in willows, sticklebacks, cichlids, frogs, and lizards, often coinciding with zones of hybridization or secondary contact^54-60^. Our findings extend this pattern to darters, positioning sex chromosome turnover as a general mechanism for the evolution of reproductive isolation.

### Mitonuclear conflict and sex-linked incompatibilities

We observed that the sex chromosomes exhibit widespread segregation distortion in *E. spectabile* x *E. caeruleum* backcross hybrids, with chromosome 23 harboring more than twice as many distorted SNPs as any other chromosome^24^. While this pattern may reflect Dobzhansky-Muller incompatibilities involving deleterious Y-linked variants, our analyses reveal that these regions are also enriched for nuclear-encoded mitochondrial genes, especially those in the OXPHOS pathway. Such enrichment raises the possibility that disrupted coevolution between mitochondrial and nuclear genomes contributes to hybrid dysfunction.

Allele frequencies at many sex-linked SNPs varied with mitochondrial haplotype (Figure 5), consistent with mitonuclear interactions. Although survival asymmetries were not detected^24^, these mismatches may affect subtler aspects of fitness such as fertility or metabolic efficiency. This aligns with the “mother’s curse” hypothesis, which predicts male-biased effects of mitochondrial mutations due to maternal inheritance^61^. Because Y-linked mitonuclear genes cannot co-inherit with mtDNA, this arrangement may constrain compensatory evolution and drive recurrent mitonuclear conflict. Such conflict could favor sex chromosome turnover as an evolutionary “escape hatch”, relocating sex determination away from incompatible loci (Fig. 4C). Our findings reported here support this model, as independently derived sex chromosomes in multiple species contain mitonuclear genes within highly sex-associated loci, and hybrid incompatibilities are concentrated at these same loci.

**Figure 5.**
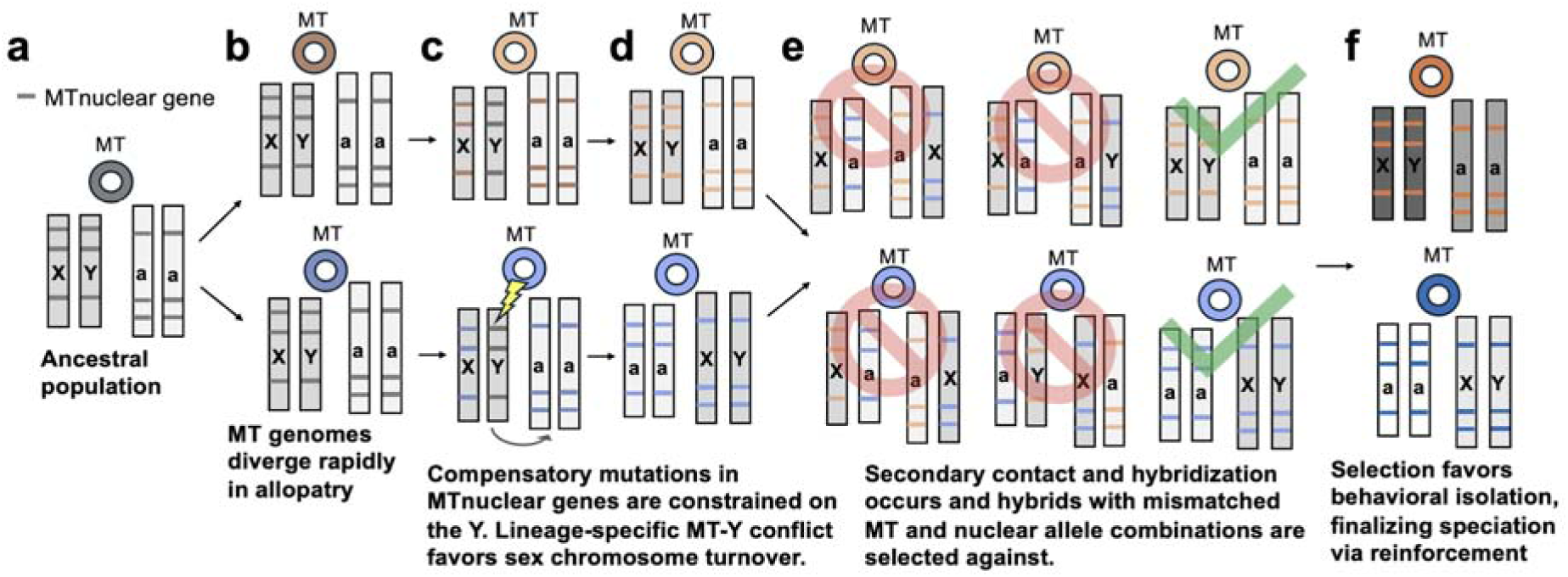
Repeated turnover of sex chromosomes as a mechanism to resolve mitonuclear conflict and promote speciation. (a) In the ancestral population, mitochondrial (MT) and nuclear genomes are co-adapted, including interactions between MT-encoded and nuclear-encoded proteins in OXPHOS complex I. (b) In allopatry, mitochondrial genomes can rapidly accumulate lineage-specific substitutions due to reduced effective population size, maternal inheritance, and relaxed purifying selection. (c) To maintain OXPHOS function, compensatory substitutions arise in nuclear-encoded mitochondrial (MTnuclear) genes. When these genes are in non-recombining regions of the Y chromosome, coevolution is constrained, exacerbating mitonuclear conflict in males. This selects for the turnover of sex-determining loci to new chromosomes, effectively unlinking sex determination from incompatible MTnuclear genes and restoring co-adaptation. (d) Each lineage accumulates distinct, co-evolved sets of mitochondrial and nuclear alleles. (e) Upon secondary contact, hybridization mixes incompatible mitochondrial and nuclear genomes. Conflict is especially pronounced when nuclear-encoded OXPHOS genes under selection reside on divergent sex chromosomes. (f) Selection acts against hybrids with mismatched mitonuclear genotypes, promoting the evolution of enhanced behavioral isolation (reinforcement) and contributing to the completion of speciation.

Previous work in darters has shown evidence of positive selection in both mitochondrial genes (e.g., *mt-nd2*, *mt-nd3*) and nuclear OXPHOS genes (e.g., *ndufs7*, *ndufb5*)^43, 44^, with many nuclear complex I genes showing derived coding changes between *E. spectabile* and *E. caeruleum* with predicted effect on protein function (Extended Data Table 3). Notably, several *E. spectabile* populations carry introgressed mitochondrial genomes from *E. caeruleum* or *E. radiosum*^62^. Given their smaller effective population size and lower nucleotide diversity, *E. spectabile* may harbor mildly deleterious mtDNA mutations, making their mitochondrial genome more prone to replacement and more likely to generate mitonuclear incompatibilities in hybrids^63^. Indeed, the enrichment of *E. caeruleum* ancestry on chromosome 23 in backcrosses to *E. spectabile* females may reflect selection for *E. caeruleum* alleles that restore mitonuclear compatibility^24^. Together, these findings suggest that ecological specialization, demographic history, and sex chromosome structure interact to shape the genomic landscape of reproductive isolation in darters. Future work linking mitonuclear variation to metabolic function will be essential to understanding the fitness consequences of this conflict.

### Conclusions

Our study reveals how sex chromosome turnover, hybrid incompatibility, and mitonuclear conflict interact to shape the genomic architecture of reproductive isolation. In darters, rapid sex chromosome evolution, the accumulation of incompatibilities on non-recombining regions, and the ecological and demographic context of mitochondrial variation combine to generate strong postzygotic barriers. As sex-linked and mitonuclear incompatibilities are increasingly recognized across taxa^3, 7, 64, 65^, darters offer a tractable model for dissecting how genomic regions shaped by sex-specific evolutionary pressures can become hotspots of reproductive isolation, revealing fundamental principles of the origin of biodiversity.

## METHODS

### Etheostoma caeruleum reference genome sequencing and assembly

We recently published a chromosome-level annotated reference genome for the orangethroat darter, *E. spectabile* (GCF_008692095.1)^24^. To account for structural and sequence changes between *E. spectabile* and *E. caeruleum* that could lead to reference-bias in our downstream analyses^66^, here we sequenced and assembled a reference genome for *E. caeruleum*. Briefly, DNA was isolated from white muscle tissue from a single male *E. caeruleum* from an allopatric population (Kalamazoo River, MI). A genomic library was prepared using a Chromium Genome Library Kit and Gel Bead Kit v2 and sequenced across two lanes of NovaSeq S4 2x150 PE (multiplexed with other 10X libraries from another project) at the University of Minnesota Genomics Center (Minneapolis, MN). Reads were assembled *de novo* using Supernova (v2.1.1)^67^ and specifying the pseudohap output to reconstruct phased scaffolds from 10X Genomics linked-read data into a single fasta formatted assembly. The assembly was further scaffolded in Chromonomer^68^ using the previously published *E. caeruleum* linkage map^24^. Scaffolds output from Chromonomer were then aligned to the orangethroat darter reference genome with RAGTAG (v2.0.1)^69^; to generate a pseudo chromosome-level assembly for *E. caeruleum*. We evaluated the completeness of the assembly with BUSCO and annotated the assembly with BRAKER3 and eggNOG-mapper (v2) (see Supplemental Materials).

### Whole genome re-sequencing

DNA was isolated from white muscle tissue of *E. spectabile*, *E. pulchellum*, *E. caeruleum*, *E. radiosum*, and *E. flabellare* using a Puregene kit (Qiagen; www.qiagen.com). We sampled species across the *Oligocephalus* subgenus, including the hybridizing pair *E. spectabile* and *E. caeruleum*, to characterize sex chromosome turnover. To test whether sex chromosome identity is conserved within the orangethroat lineage, we included *E. pulchellum* (a close relative of *E. spectabile* within the orangethroat darter complex)^70^. We further included *E. radiosum*, which represents another lineage within *Oligocephalus* that hybridizes with *E. pulchellum*^71^. *E. flabellare*, a distantly related species outside of *Oligocephalus*, was included as an outgroup for phylogenomic and introgression analyses (TWISST, *fdM*). Collection locations and sample sizes are provided in Extended Data Table 1. DNA quantification and checks for sample purity were performed on a NanoDrop 1000 spectrophotometer. All samples were stored in -20° C until library preparation and sequencing at the Texas A&M AgriLife Genomics and Bioinformatics Service (TxGen; College Station, TX). DNA quality control testing was done on a DropQuant spectrophotometer to obtain OD ratios, read by picogreen to obtain concentrations, diluted to 5ng/µL, and run on the Agilent Fragment Analyzer using a high sensitivity genomic DNA kit. Libraries were generated using NEXTFLEX Rapid XP DNA-Seq Kits and automated on a PerkinElmer robotics system. Resulting libraries were sequenced across five lanes on a NovaSeq X Plus 25B flowcell (4,875 Gb total sequencing) to a mean depth of 27X coverage of the approximately 1 Gb genome.

We used FastQC (v0.11.8) to check the quality of the resulting raw reads and to generate sample-specific read count statistics. We trimmed adapters using Cutadapt (v4.2) with barcodes specified for each individual and performed quality trimming using Trimmomatic (v0.39). We required a minimum quality score of 30 across a 6□bp sliding window and discarded reads with a length of <40 nucleotides. After merging pair end reads with seqtk (v1.4), we used bwa (v0.7.17) to align reads for each sample to both the published *E. spectabile* reference genome (GCF_008692095.1)^24^and our *de novo E. caeruluem* reference genome. We used Picard (v2.18.27) to remove duplicates and add read group information and used samtools (v1.20) to split de-duplicated bams into mapped and unmapped reads.

### Genotype Calling

We conducted genotype calling on mapped reads using the Genome Analysis Tool Kit (GATK) following the GATK Best Practices guidelines^72-74^. First, we used the HaplotypeCaller tool in GATK (v4.5) to generate per-individual gvcfs from the mapped bam files. Then, we used the GenotypeGVCFs tool in GATK (v4.5) to produce vcf files for each chromosome and all unplaced scaffolds. We applied hard filters using the SelectVariants and VariantFiltration tools in GATK (v4.5). To do so, we subset the VCFs for each chromosome and unplaced scaffold into invariant, SNP, and mixed/indel sites and applied filters separately following GATK best practices. We finally used the MergeVcfs tool in GATK (v4.5) to re-combine all subset VCFs for each chromosome and unplaced scaffold.

After genotype calling, we took several steps to ensure the quality of our data. We used VCFtools (v0.1.16) to remove sites with greater than 20% missing data in each population. We used the ‘vcftools -- exclude-bed’ option to remove repetitive regions identified by RepeatMasker with files downloaded from NCBI (GCF_008692095.1). Indels and 10 bp immediately upstream and downstream around each indel were removed using a custom python script. We calculated heterozygosity at each site for each population in R (v4.0.3) using the vcfR package^75^. We excluded sites where every sample in a given population was heterozygous, which is indicative of collapsed paralogs. We additionally used BCFtools (v1.19) to only retain biallelic SNPs and remove variants with a minor allele frequency <1%.

### Sex association tests

We performed a genome-wide association study (GWAS) to identify genomic regions associated with sex using linear mixed models in GEMMA (v0.98.5)^76^. Separate analyses were conducted on *E. spectabile*, *E. pulchellum*, *E. caeruleum*, and *E. radiosum*. For each species, we first estimated a centered relatedness matrix using the ‘-gk 2’ option from PLINK-formatted genotype data to account for population structure and relatedness among individuals. The resulting matrix was used as input for a linear mixed model association test, using the ‘-lmm 4’ option, which computes the Wald, likelihood ratio, and score test statistics for each SNP.

We analyzed a total of 3,550,084 SNPs for a pooled set of allopatric and sympatric *E. spectabile* aligned to the *E. spectabile* reference genome. Additionally, we analyzed a total of 4,627,034 SNPs for a pooled set of allopatric and sympatric *E. caeruleum* aligned to the 10X *E. caeruleum* reference genome assembly. A total of 10,747,161 SNPs were included in the *E. radiosum* analysis and 3,701,092 SNPs were included in the *E. pulchellum* analysis (both aligned to the *E. spectabile* reference genome). For *E. spectabile* and *E. caeruleum*, we also analyzed allopatric and sympatric populations separately for each species to assess whether sex-associated regions are conserved among populations within species. We visualized the results of our genome-wide association study (GWAS) using Manhattan plots generated in R with the qqman^77^ and ggplot2^78^ libraries.

As a second approach to identify sex-associated sites that show heightened differentiation between males and females, we calculated intersex Weir’s F_ST_ with VCFtools (v0.1.16)^79^ at each site across the genome in both allopatric and sympatric populations of *E. spectabile* and *E. caeruleum*. We calculated intersex F_ST_ for each gene in the species-specific annotation and identified outliers using a custom Python script. We identified outlier genes with exceptionally high genetic differentiation (>95th percentile) between males and females in each population.

### Investigating heterogamety

We took several complementary approaches to investigate which sex may be heterogametic in darters. Although the sex determining system has not previously been investigated in any other darter species, male heterogametic sex determining systems have been identified in several non-darter percids^80^, which are estimated to have the most recent common ancestor with darters around 58-66 million years ago^81^. For a species with male heterogametic sex determination, males are expected to be completely heterozygous (one X and one Y allele) at the sex determining region. Additionally, females would be expected to be homozygous (two X alleles) across the sex determining region. For *E. spectabile*, *E. caeruleum*, and *E. radiosum*, we calculated the difference between male and female observed heterozygosity at each site across the genome using the vcfR package in R v4.0.3^75^. To reduce noise and computational burden, we aggregated sites into 250 kb non-overlapping windows, calculating the mean male – female heterozygosity within each bin. For each species, bins were assigned as either “sex chromosome” (if they mapped to the chromosome previously identified as sex-linked) or “autosome” (all other chromosomes). We then compared window means between sex chromosome and autosomal bins using a Wilcoxon rank-sum test. We additionally calculated Cliff’s delta as a nonparametric measure of effect size. To quantify uncertainty in the mean difference between groups, we estimated a 95% confidence interval using the parametric standard error of the difference in means. If the sex chromosomes display significantly male-biased heterozygosity, this would support the hypothesis that the species has a male heterogametic sex determining system. The converse would support a female heterogametic system.

The sex that is heterogametic is also predicted to accumulate sex-specific SNPs on the Y or W. To test this, we calculated sex differences in SNP density across each chromosome in *E. spectabile*, *E. pulchellum*, *E. caeruleum*, and *E. radiosum*. We filtered the dataset to include only biallelic SNPs from males and females aligned to the same-species reference genome, retaining variants with a quality score >30 and minimum coverage of 20^81,82^. Using BEDtools, we quantified the number of SNPs in 10 kb windows across the genome for each sample. Following Darolti et al. ^82^, male-biased SNP density was calculated as log_2_(average male SNP density) - log_2_(average female SNP density). We applied Wilcoxon rank sum tests to compare male-biased SNP density on the sex chromosome versus the autosomes. Together, these analyses of male-biased SNP density and heterozygosity were used to confirm and refine the boundaries of candidate sex-determining regions indicated by sex association tests.

### Genetic mechanisms for male-biased F1 hybrid clutches

Our preliminary genomic analyses suggested distinct sex chromosomes in *E. spectabile* and *E. caeruleum*, with potential support for a neo-sex chromosome in *E. caeruleum* and maintenance of XY male heterogamety in both species. We previously found that laboratory-generated F1 hybrid crosses in both directions result in clutches with dramatically male-skewed sex ratios (♀ *E. spectabile* x ♂ *E. caeruleum*: mean ± SE prop male = 0.84 ± 0.10; ♀ *E. caeruleum* x ♂ *E. spectabile*: mean ± SE prop male = 0.95 ± 0.04)^28^. Notably, the number of surviving offspring in F1 and parental crosses do not differ, suggesting genetically female individuals are not missing from F1 crosses, but are phenotypically male^28^.

To explain these striking male-skewed sex ratios in F1 hybrids, we next considered what type of sex chromosome transition could account for the pattern. Our genomic analyses indicate that the transition in sex chromosome system that occurred between *E. spectabile* and *E. caeruleum* was (1) heterologous, involving a change in which chromosome pair serves as the sex chromosomes, and (2) maintained male heterogamety. However, even given these parameters, there are several different evolutionary pathways through which the sex chromosome system can evolve. A key distinction among these pathways is whether the transition occurred via the invasion of a ‘dominant’ or ‘recessive’ new sex-determining mutation. A ‘dominant’ sex determiner indicates that one copy of the sex-determining mutation is sufficient to determine sex (e.g., as SRY on the Y chromosome behaves in mammals), while a ‘recessive’ sex determiner indicates that two copies of the sex-determining mutation are required to determine sex (e.g., as observed in the ‘counting X’s’ system found in *Drosophila melanogaster*)^31,83^.

To clarify the nature of the sex chromosome transition observed here, we calculated the expected sex ratios that would arise if *E. caeruleum* had undergone different types of sex chromosome transitions from the ancestral *E. spectabile* system. Based on these results, and on data from the candidate gene analysis described below, we can predict the likely evolutionary pathway through which the novel sex chromosome system arose in *E. caeruleum*.

### Candidate genes for sex determination

We sought to identify candidate sex-determining genes located within peaks of the genome-wide association study plots. Using the *E. spectabile* annotation file available on NCBI, we applied bedtools intersect to extract genes within 50 kb upstream or downstream of GEMMA outlier SNPs in *E. spectabile* and *E. radiosum*. *E. pulchellum* was not included in this analysis, as preliminary results indicated concordance in sex-linked regions with *E. spectabile* (a close relative within the orangethroat darter complex). For *E. caeruleum*, we used the annotation file generated from our 10X genome assembly to identify genes within 50 kb of outlier SNPs. We then conducted separate gene ontology (GO) analyses for each species-specific gene list to evaluate their potential functions.

Preliminary analyses of outlier genes indicated that one significantly sex-associated region for *E. caeruleum* was within a long non-coding (lnc) RNA. To identify potential gene targets for this lncRNA, we conducted a blastN search and retained gene hits that had at least 85% percent identity and an E-value <0.00001. We conducted a separate GO enrichment analysis on this set of putative target genes to infer function of the lncRNA.

### Ancestry calling and introgression analyses

Theory predicts that regions of the genome with reduced recombination within species (such as sex determining loci) are likely to be resistant to introgression and show a depletion of minor parent ancestry relative to the genomic background^84-86^. We used topology weighting and calculated ABBA-BABA statistics across the genome to identify regions that remain highly divergent and resistant to introgression between *E. spectabile* and *E. caeruleum* in the face of ongoing gene flow in sympatry (providing candidate regions underlying reproductive isolating barriers) and to ask whether patterns differ between sex chromosomes and autosomes (see Supplementary Methods).

### Mapping mitonuclear associations

Preliminary sex determination candidate gene analysis revealed an enrichment of mitochondrial-associated genes in highly sex-linked regions within *E. spectabile*, *E. caeruleum*, and *E. radiosum*. We re-analyzed previously published genotype data for lab-generated backcross hybrids between *E. spectabile* and *E. caeruleum*^24^ to test for an association between the mitochondrial haplotype (*E. spectabile* or *E. caeruleum*) and ancestry across the nuclear genome. Backcrosses were created in both possible directions by crossing *E. spectabile* x *E. caeruleum* F1 hybrid males to non-admixed females of both parental species. Offspring surviving to three days post-hatching (n=13 *E. caeruleum* backcross fry, n = 36 *E. spectabile* backcross fry) were genotyped with RADseq^24^. A total of 16,585 SNPs present in the backcross RADseq data were differentially fixed in the parental species and we retained 8,177 SNPs after pruning for linkage.

To identify nuclear loci exhibiting non-random associations with mitochondrial background, we performed a chi-square test of independence for each SNP. The test compared observed nuclear allele frequencies between individuals carrying *E. spectabile* versus *E. caeruleum* mitochondrial haplotypes, based on allele counts in backcross hybrid offspring. For each SNP, we constructed a 2×2 contingency table representing counts of the minor and major alleles in individuals with each mitochondrial haplotype. We then used chi square tests to assess whether nuclear allele frequency was independent of mitochondrial haplotype. Observed and expected minor allele counts were calculated using the number of chromosomes sampled and the expected frequency assuming no association between nuclear genotype and mitochondrial background. To ask whether any individual chromosomes were enriched for mitonuclear associations, we used Fisher’s exact tests and applied an FDR correction for multiple testing.

To further investigate whether the *E. spectabile* and *E. caeruleum* sex chromosomes (9 and 23, respectively) exhibited non-random shifts in allele frequency in backcross hybrids associated with mitochondrial background, we calculated the difference in minor allele frequency (ΔMAF) at ancestry-informative sites between offspring carrying orangethroat darter mitochondria and those carrying *E. caeruleum* mitochondria at each site. We then tested whether the mean ΔMAF on chromosomes 9 and 23 differed significantly from genome-wide expectations using two complementary approaches. First, for each chromosome, we randomly sampled the same number of ΔMAF values from the rest of the genome (excluding chromosomes 9 and 23), repeated this process 10,000 times, and compared the observed chromosome-specific mean ΔMAF to the null distribution. Second, we used a Wilcoxon rank-sum test to compare the distribution of ΔMAF values on chromosomes 9 and 23 against the distribution from all other chromosomes. We used bedtools (v2.3.1)^87^ to extract all genes from the *E. spectabile* genome annotation occurring within 50 kb from SNPs that had significant p-values in our association mapping between nuclear genotype and mitochondrial background. We used GO analysis (https://geneontology.org/) to infer if these genes were enriched for any set of biological processes.

To assess whether mitonuclear genes are non-randomly distributed across the genome, we tested for chromosomal enrichment or depletion after accounting for differences in chromosome length. Out of 1,159 nuclear encoded genes with known mitochondrial-related functions present in the MitoCarta2.0 database^88^, 867 orthologs are present in the *E. spectabile* reference genome annotation (Supplementary Table 1). Mitonuclear gene density on each chromosome was calculated as the number of mitonuclear genes per Mb. We calculated the expected number of mitonuclear genes per chromosome under the null hypothesis that genes are distributed proportionally to chromosome length. Z-scores were computed for each chromosome to quantify the deviation of observed counts from expected values. To account for multiple testing across the 24 chromosomes, we applied a Benjamini-Hochberg FDR correction.

### Functional implications of mitonuclear substitutions

To identify potentially deleterious substitutions contributing to mitonuclear incompatibilities between *E. spectabile* and *E. caeruleum*, we examined the predicted effect of derived variants present in nuclear-encoded and mitochondrial-encoded genes in complex I of the oxidative phosphorylation (OXPHOS) pathway. Specifically, we focused on five nuclear-encoded genes located on the sex chromosomes (*ndufa7*, *ndufa11*, *ndufa12*, *ndufb5*, and *ndufs7*) and three mitochondrial-encoded genes that contain variants previously shown to be under positive selection in these species (*mt-nd2*, *mt-nd3*, *mt-nd5*). Using our annotated *E. caeruleum* reference genome, we extracted coding sequences for these genes. We then aligned the sequences with homologous genes from other darter and percid species for which annotated protein sequences were available on NCBI.

Protein alignments were performed using Clustal Omega (v1.2.4) to identify sites with derived amino acid substitutions that were unique to *E. spectabile* and/or *E. caeruleum*. The functional effects of nonsynonymous substitutions were predicted using the SIFT algorithm (Sorting Intolerant From Tolerant), which assesses whether an amino acid substitution is likely to affect protein function based on sequence conservation and physicochemical properties^89^.

To evaluate the potential structural consequences of the identified substitutions, we downloaded the crystal structure of bovine mitochondrial complex I (PDB: 5XTD) and visualized orthologous residue positions using PyMOL (v3.0.5). Structural homology was inferred based on known alignments of complex I subunits, allowing visualization of residue positions within the multi-subunit architecture of the complex. Residues predicted to be deleterious by SIFT and uniquely derived in *E. spectabile* or *E. caeruleum* were annotated and highlighted for further interpretation in the context of hybrid incompatibility.

## DATA AVAILABILITY

All raw reads have been deposited to NCBI SRA (SUB14878521) and will be released upon publication.

## CODE AVAILABILITY

All scripts necessary to recreate data processing and analysis are available at https://github.com/rachelmoran28.

## ACKNOWLEDGEMENTS

The treatment of animals used in this study was in compliance with Texas A&M University’s Institutional Animal Care and Use Committee (IACUC) under AUP #2023-0007. We thank the University of Minnesota Genomics Center and TxGen Genomics Core for their guidance and performing the 10X library preparations and Illumina sequencing. The Texas A&M University HPRC provided resources that contributed to the research results reported within this paper. Funding was supported by NSF IOS #2338043 to R.L.M., an AGA EECG to R.L.M., and a Texas A&M University Merit Fellowship to W.V.R. We are grateful to Taylor N. Black, Kiedon Bryant, and Brynn Johnson for assistance in the field. Fish were collected under Illinois Department of Natural Resources permit A24.6515, Michigan Department of Natural Resources permit FSCP02212024085151, Texas Parks and Wildlife Department permit SPR-0323-023, Oklahoma Department of Wildlife Conservation license 10775512, and Arkansas Game and Fish Commission permit 20620232 to RLM.

## AUTHOR CONTRIBUTIONS

W.V.R. analysed the data and drafted the manuscript. M.D. assisted with data analysis and manuscript preparation. D.K. contributed to data analysis. P.M. conducted analyses and contributed to manuscript writing. R.L.M. conceived and designed the project with M.D. and W.V.R., and analysed data. All authors contributed to writing and revising the manuscript.

Supplementary Information is available for this paper.

Correspondence and requests for materials should be addressed to Rachel Moran.

---

**Extended Data Figure 1.**
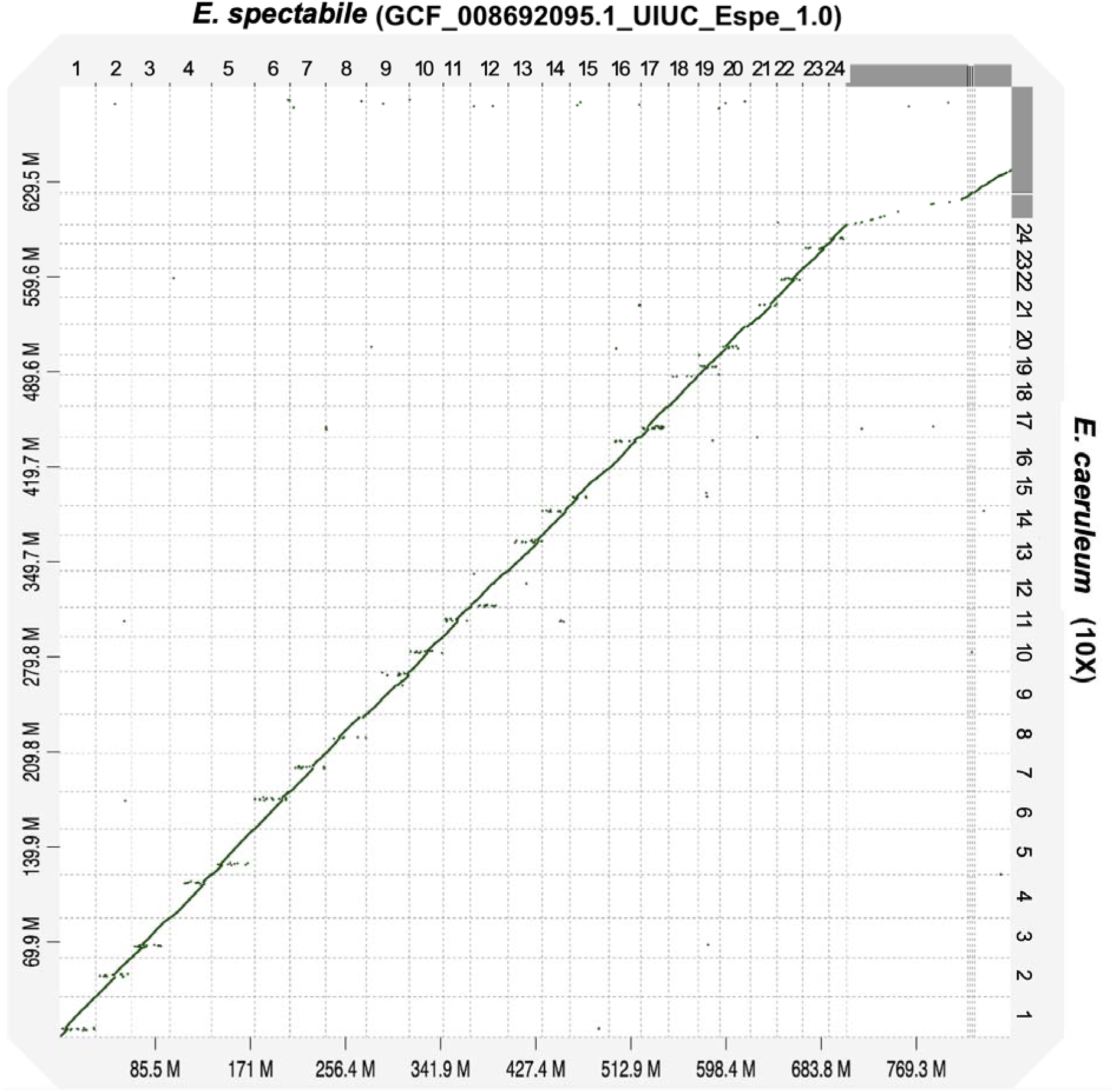
High degree of synteny between *E. spectabile* and *E. caeruleum* genome assemblies. A D-Genies dot plot^90^ between the orangethroat darter (*Etheostoma spectabile*) reference genome (x-axis) and our de novo 10X rainbow darter (*Etheostoma caeruleum*) reference genome (y-axis). Each dot represents a sequence match between genomes, with the diagonal line indicating synteny. Darker green indicates a higher percent identity match and lighter orange indicates a lower percent identity match. Off-diagonal points may reflect structural variation, rearrangements, or mapping artifacts. The dense diagonal signal suggests high collinearity between the *E. spectabile* sympatric individuals and the reference genome.

**Extended Data Figure 2.**
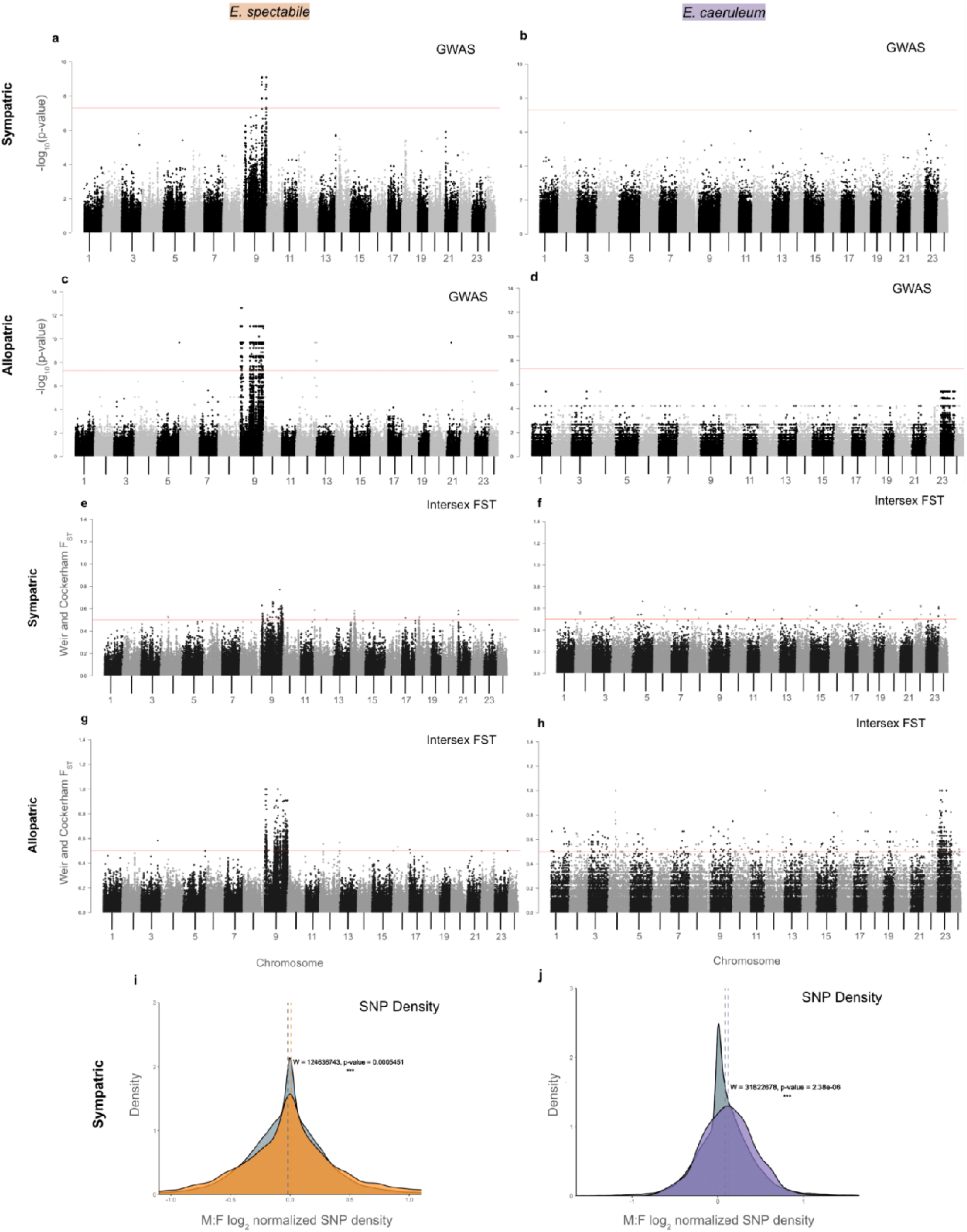
Sex chromosome incompatibilities diverged prior to secondary contact. Genome-wide association study using sex as a phenotype for sympatric-only populations of (a) *E. spectabile* (orangethroat darters) and (b) *E. caeruleum* (rainbow darters) indicated a weaker signal of sex association than in the allopatric populations of the same species (c, d). A total of 918,863 single nucleotide polymorphisms (SNPs) were analyzed for sympatric *E. spectabile* and 4,056,249 SNPs were analyzed for the sympatric *E. caeruleum*. Manhattan plots displaying genome-wide Weir and Cockerham FST values for (e) sympatric orangethroat, (f) sympatric rainbow, (g) allopatric orangethroat, and (h) allopatric rainbow samples. Chromosome 9 still has a relatively strong signal in sympatry for orangethroat darters, but there is no significant signal across the genome of sympatric rainbow darters, suggesting the signal weakens in sympatry. Density plots showing normalized log_2_ difference between male and female SNP density for (i) sympatric orangethroat darters aligned to the orangethroat reference genome and (j) sympatric rainbow darters aligned to the rainbow darter reference genome. In (i), the sex chromosome (chromosome 9) is shown in orange. In (j), the sex chromosome (chromosome 23) is shown in purple. Autosomes are shown in gray. In both (i) and (j), sex chromosomes have significantly higher male-biased SNP density compared to autosomes. Male-skewed SNP density is the only supporting evidence for sex chromosome identity that has a significant signal for both species in sympatry.

**Extended Data Figure 3.**
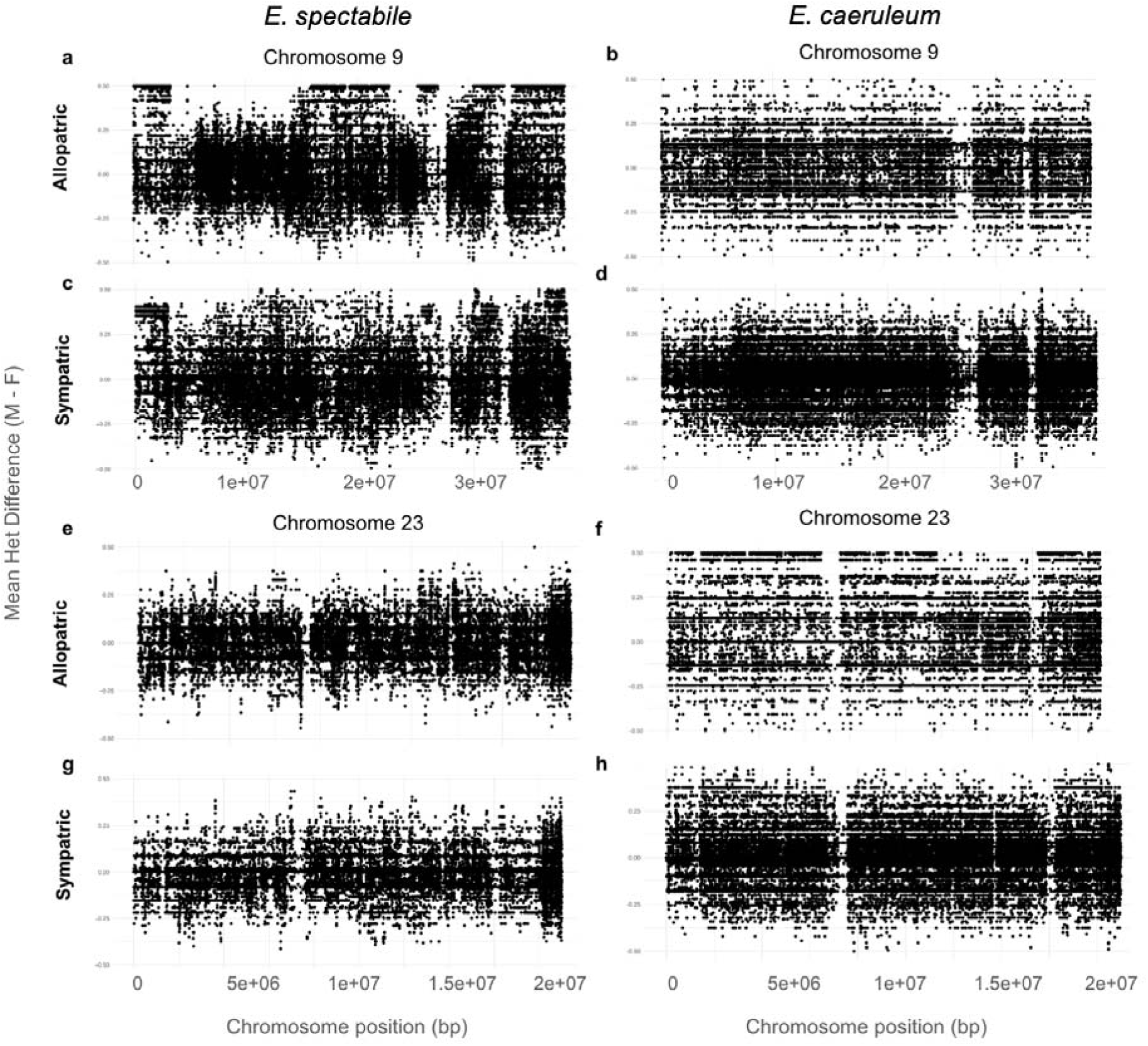
Male-biased heterozygosity on putative sex chromosomes. Mean difference in heterozygosity between males and females across chromosomes 9 and 23 for *E. spectabile* and *E. caeruleum*. Data are shown for allopatric populations of *E. spectabile* (Sangamon River, IL) (a,e) and *E. caeruleum* (Kalamazoo River, MI) (b,f) as well as a sympatric population of the two species (Vermilion River, IL) (c,d,g,h). Heterozygosity is strongly biased towards males on the putative sex chromosome (a,c) but not on the autosomal chromosome (e,g) in the *E. spectabile* comparison, indicating a male heterogametic XY system. A similar pattern emerges for the rainbow darter populations, with heterozygosity being strongly biased towards males on the putative sex chromosome in allopatry (f) but not on the autosomal chromosome (b), indicating a male heterogametic XY system. Sympatric *E. caeruleum* show a decreased male-biased heterozygosity across the putative sex chromosome (h), likely due to gene flow with sympatric *E. spectabile*.

**Extended Data Figure 4.**
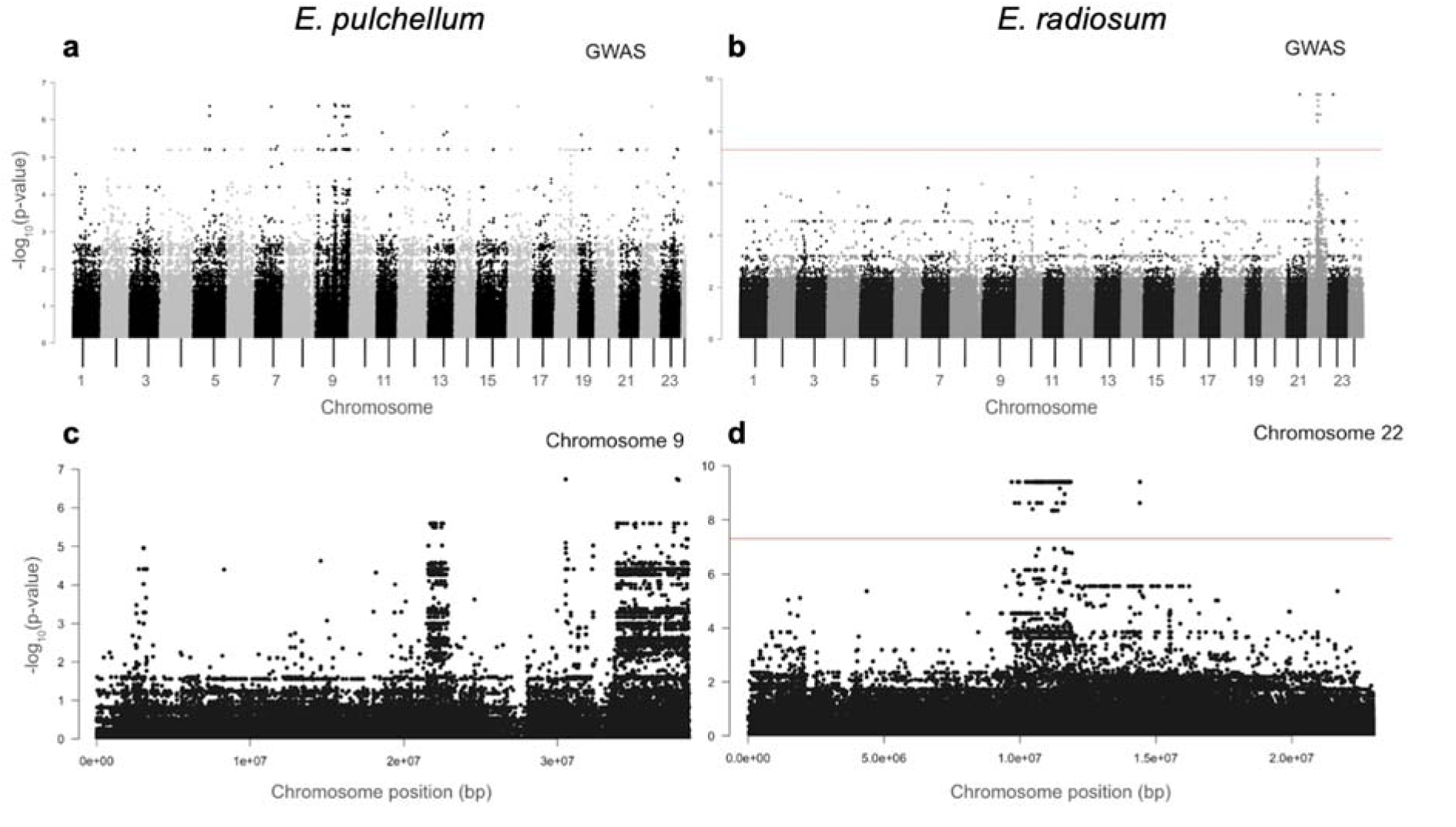
Sex chromosome identity in *E. pulchellum* and *E. radiosum.* Genome-wide association study on sex showed peaks on chromosome 9 in *E. pulchellum* (a member of the orangethroat darter species complex) (a), and on chromosome 22 in orangebelly darters, *E. radiosum* (b). A total of 10,747,161 SNPs (n=10 males, n=16 females) were included in the *E. radiosum* analysis and 3,701,092 SNPs (n=13 males, n=13 females) were included in the *E. pulchellum* analysis. Chromosome-level GWAS using sex as a phenotype subset to only chromosome 9 in *E. pulchellum* (c) and chromosome 22 in *E. radiosum* (d).

**Extended Data Figure 5.**
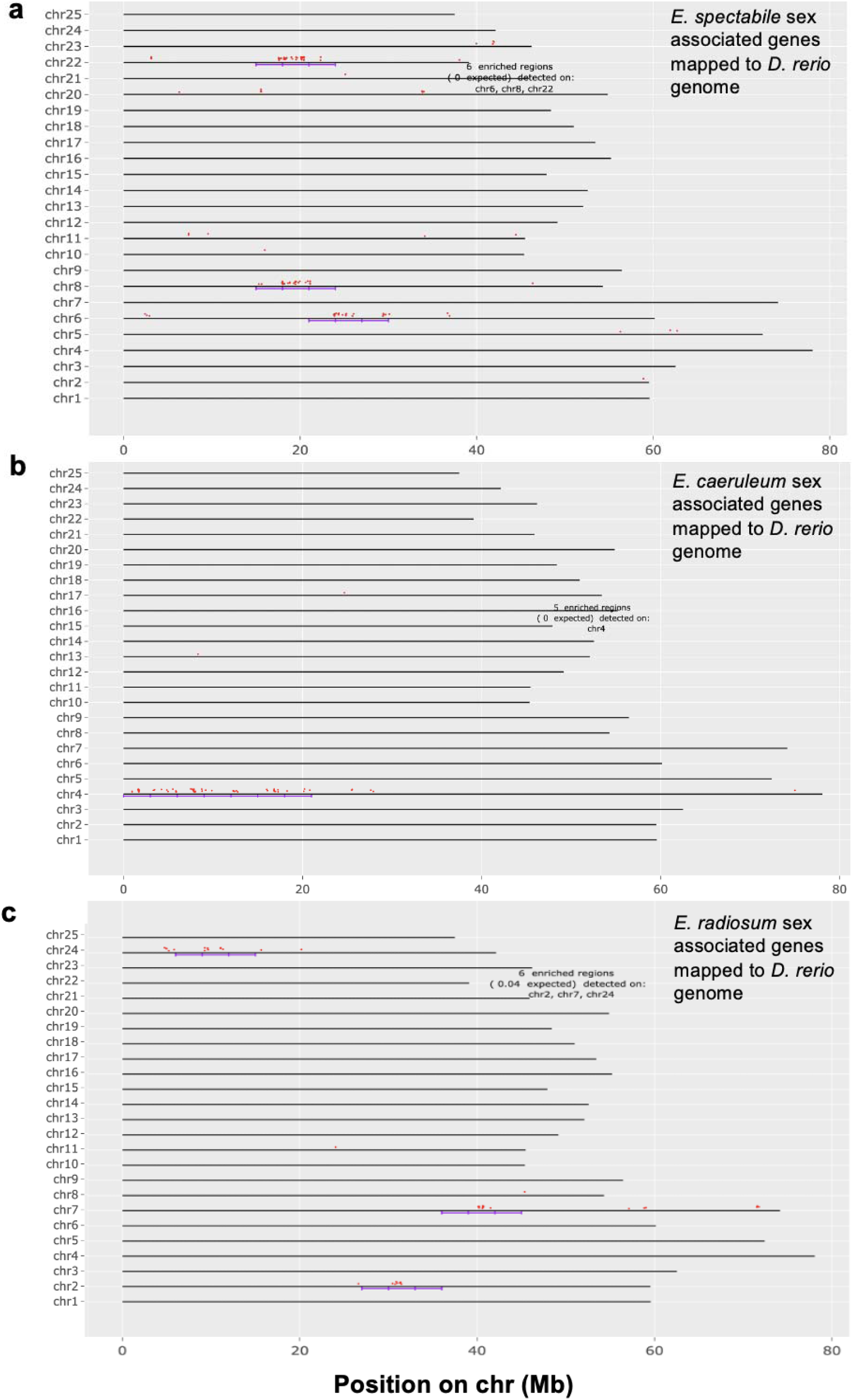
Synteny mapping indicates sex associated regions are shared between *Etheostoma caeruleum* and *Danio rerio*. Genomic distribution of sex-determining candidate genes from (a) *E. spectabile* chromosome 9, (b) *E. caeruleum* chromosome 23, and (c) *E. radiosum* chromosome 22 mapped to the zebrafish (*Danio rerio*) genome using ShinyGO v0.82. Each horizontal black line represents a zebrafish chromosome, with red dots indicating darter candidate genes (Supplementary Table 4). The genome was scanned using a sliding window approach (window size = 6 Mb, 2 steps per window), and enrichment of candidate genes within each window was assessed using a hypergeometric test with an FDR cutoff of 1e-05. Purple brackets indicate windows significantly enriched for candidate genes compared to the background gene density. Notably, the right arm of chromosome 4 (Chr4R) in zebrafish is sex-linked and also harbors a cluster of four enriched windows of rainbow darter sex-associated genes (with 0 expected by chance), suggesting a region homologous to rainbow darter chromosome 23 that may play a conserved role in sex determination.

**Extended Data Figure 6.**
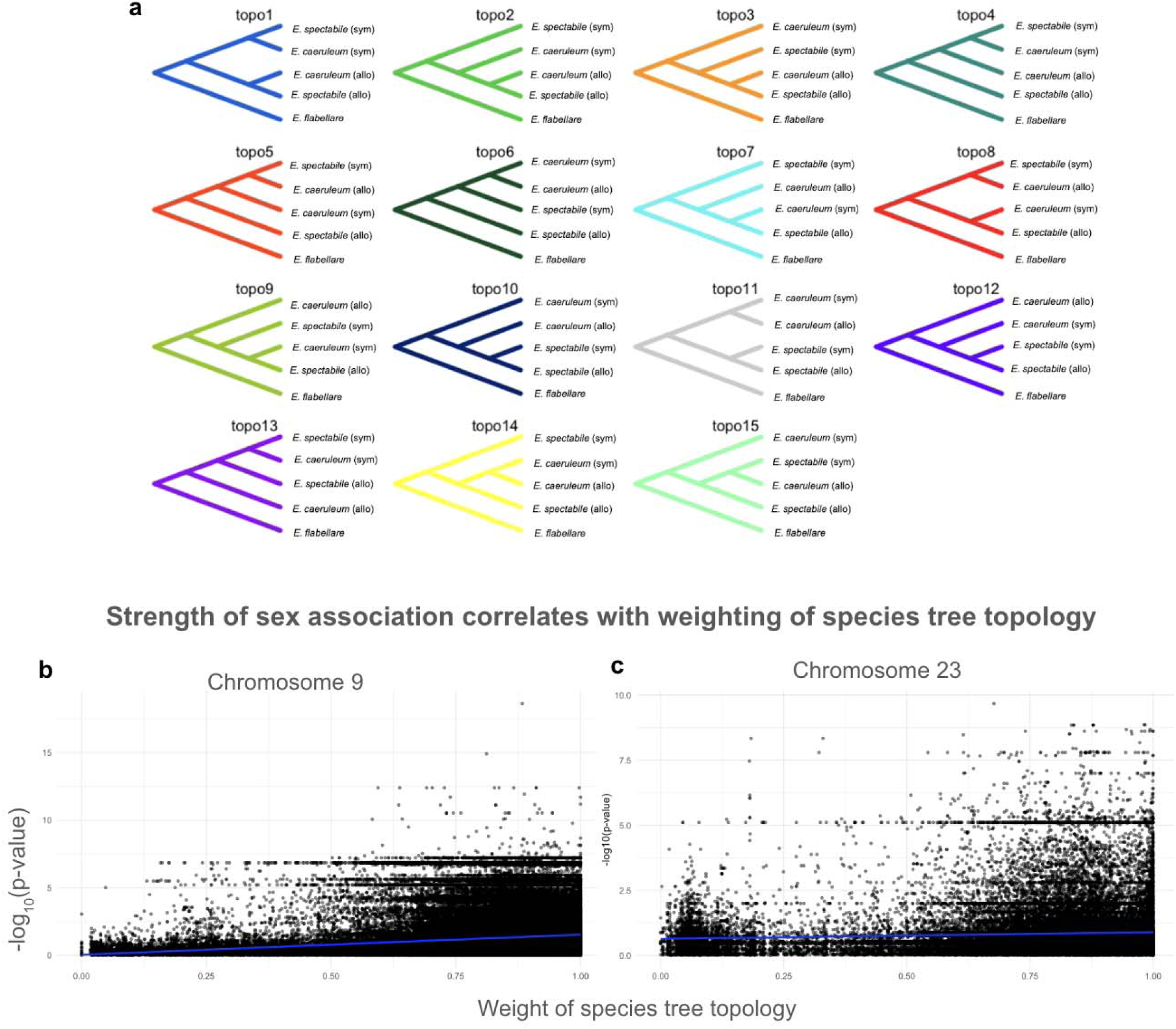
Sex-linked regions show reduced phylogenetic discordance, implying less introgression. (a) All possible topologies inferred by TWISST, a method that tracks the support for different phylogenetic relationships across the genome. We included one allopatric (allo) and one sympatric (sym) population each of *E. caeruleum* and *E. spectabile*, and *E. flabellare* served as an outgroup. Each panel represents a different way the five populations could be related based on local genomic windows. Topology 11 (topo11), shown here in grey, reflects the expected species tree (indicate no introgression between sympatric orangethroat and rainbow darters). (b, c) TWISST analyses revealed that genomic regions with strong sex associations (GWAS, p < 5 × 10^−8^) also had higher support for the species tree topology (a, topo11), indicating reduced phylogenetic discordance and gene flow in sex-linked regions within species. (b) *E. spectabile* chromosome 9 (r = 0.13, p < 2.2e^−16^). (c) *E. caeruleum* chromosome 23 (r = 0.06, p < 2.2e^−16^).

**Extended Data Figure 7.**
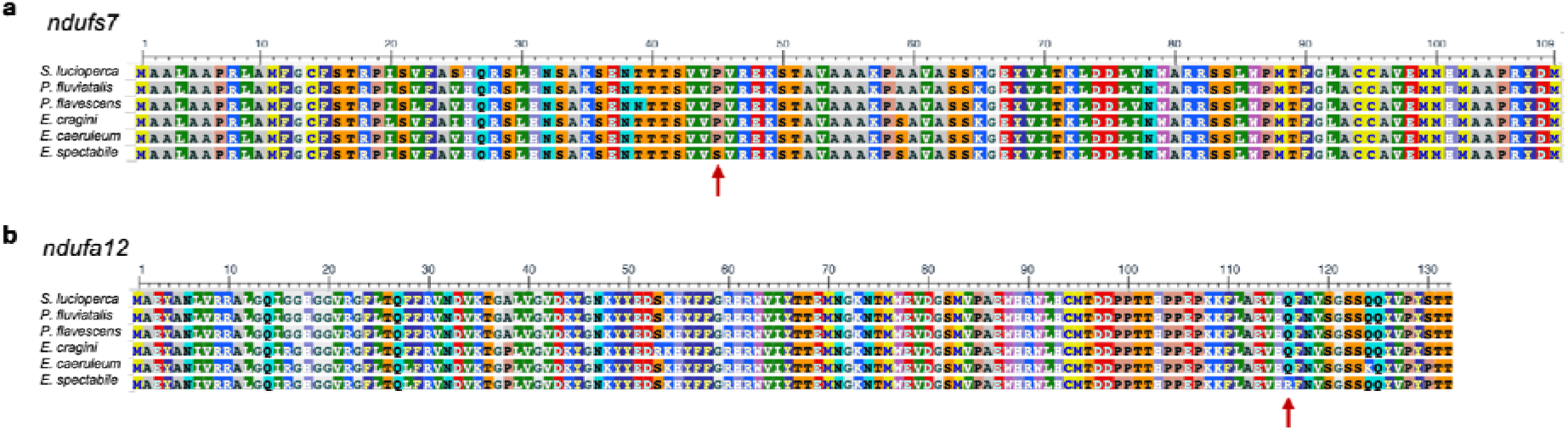
Mutations in mitonuclear proteins with high sex association impact protein function. Derived mutations predicted by SIFT to be deleterious to protein function are present in *E. spectabile* in (a) *ndufs7* (chromosome 9; SIFT score = 0.04) and (b) *ndufa12* (chromosome 23; SIFT score = 0.02).

**Extended Data Table 1.** Collection locations and sample sizes of darters used in study.

| SPECIES | SEX | COORDINATES | RIVER DRAINAGE | SAMPLE SIZE |
| --- | --- | --- | --- | --- |
| *Etheostoma caeruleum* | Male | 42.426474, -85.428319 | Kalamazoo River, MI | n = 7 |
| *Etheostoma caeruleum* | Female | 42.426474, -85.428319 | Kalamazoo River, MI | n = 7 |
| *Etheostoma caeruleum* | Male | 40.055556, -88.091689 | **Vermilion River, IL** | n = 12 |
| *Etheostoma caeruleum* | Female | 40.055556, -88.091689 | **Vermilion River, IL** | n = 12 |
| *Etheostoma spectabile* | Male | 40.027060, -88.574980 | Sangamon River, IL | n = 12 |
| *Etheostoma spectabile* | Female | 40.027060, -88.574980 | Sangamon River, IL | n = 12 |
| *Etheostoma spectabile* | Male | 40.055556, -88.091689 | **Vermilion River, IL** | n = 12 |
| *Etheostoma spectabile* | Female | 40.055556, -88.091689 | **Vermilion River, IL** | n = 12 |
| *Etheostoma pulchellum* | Male | 30.945651, -97.533463 | Brazos River, TX | n = 5 |
| *Etheostoma pulchellum* | Female | 30.945651, -97.533463 | Brazos River, TX | n = 5 |
| *Etheostoma pulchellum* | Male | 34.351361, -96.710306 | **Washita River, OK** | n = 5 |
| *Etheostoma pulchellum* | Female | 34.351361, -96.710306 | **Washita River, OK** | n = 5 |
| *Etheostoma pulchellum* | Male | 34.454389, -96.635806 | **Blue River, OK** | n = 3 |
| *Etheostoma pulchellum* | Female | 34.454389, -96.635806 | **Blue River, OK** | n = 3 |
| *Etheostoma radiosum radiosum* | Male | 34.414424, -93.630726 | Caddo River, AR | n = 5 |
| *Etheostoma radiosum radiosum* | Female | 34.414424, -93.630726 | Caddo River, AR | n = 5 |
| *Etheostoma radiosum paludosum* | Male | 34.031389, -96.728472 | **Washita River, OK** | n = 1 |
| *Etheostoma radiosum paludosum* | Female | 34.031389, -96.728472 | **Washita River, OK** | n = 4 |
| *Etheostoma radiosum cyanorum* | Male | 34.454389, -96.635806 | **Blue River, OK** | n = 4 |
| *Etheostoma radiosum cyanorum* | Female | 34.454389, -96.635806 | **Blue River, OK** | n = 7 |
| *Etheostoma flabellare* | Male | 40.027060, -88.574980 | Sangamon River, IL | n = 4 |
| *Etheostoma flabellare* | Female | 40.027060, -88.574980 | Sangamon River, IL | n = 5 |

Species name, coordinates of and formal name for the river drainages sampled, and sample size for all male and females collected for sequencing in this study. Sympatric river drainages containing two species are bolded (Vermilion River, IL = sympatry between *E. spectabile* and *E. caeruleum*; Blue River and Washita River, OK = sympatry between *E. pulchellum* and *E. radiosum* spp.

**Extended Data Table 2.** Sex ratio frequencies in heterologous transition model.

| Female genotypes |  |  |  | Male genotypes |  |  |
| --- | --- | --- | --- | --- | --- | --- |
| F1 | XX | aa |  | XY | aa | M1 |
| F2 | XX | a*a |  | XY | a*a | M2 |
| F3 | XX | a*a* |  |  |  |  |
| F4 | XY | a*a* |  |  |  |  |
| F5 | YY | a*a* |  | YY | a*a | M3 |

| CROSS | OFFSPRING PROPORTION | OFFSPRING GENOTYPE | OFFSPRING SEX | SEX RATIO |  |
| --- | --- | --- | --- | --- | --- |
| F1 x M1 | 0.5 | XX aa | F | 0.5 F | NOTE: recurrent genotype pair in orangethroat darters |
|  | 0.5 | XY aa | M | 0.5 M |  |
| F1 x M2 | 0.25 | XX aa | F | 0.5 F |  |
|  | 0.25 | XY aa | M | 0.5 M |  |
|  | 0.25 | XX a*a | F |  |  |
|  | 0.25 | XY a*a | M |  |  |
| F1 x M3 | 0.5 | XY aa | M | 1 M |  |
|  | 0.5 | XY a*a | M |  |  |
| F2 x M1 | 0.25 | XX aa | F | 0.5 F |  |
|  | 0.25 | XY aa | M | 0.5 M |  |
|  | 0.25 | XX a*a | F |  |  |
|  | 0.25 | XY a*a | M |  |  |
| F2 x M2 | 0.125 | XX aa | F | 0.625 F |  |
|  | 0.25 | XX a*a | F | 0.375 M |  |
|  | 0.125 | XX a*a* | F |  |  |
|  | 0.125 | XY aa | M |  |  |
|  | 0.25 | XY a*a | M |  |  |
|  | 0.125 | XY a*a* | F |  |  |
| F2 x M3 | 0.25 | XY aa | M | 0.25 F |  |
|  | 0.5 | XY a*a | M | 0.75 M |  |
|  | 0.25 | XY a*a* | F |  |  |
| F3 x M1 | 0.5 | XX a*a | F | 0.5 F |  |
|  | 0.5 | XY a*a | M | 0.5 M |  |
| F3 x M2 | 0.25 | XX a*a | F | 0.75 F |  |
|  | 0.25 | XY a*a | M | 0.25 M |  |
|  | 0.25 | XX a*a* | F |  |  |
|  | 0.25 | XY a*a* | F |  |  |
| F3 x M3 | 0.5 | XY a*a | M | 0.5 F |  |
|  | 0.5 | XY a*a* | F | 0.5 M |  |
| F4 x M1 | 0.25 | XX a*a | F | 0.25 F |  |
|  | 0.5 | XY a*a | M | 0.75 M |  |
|  | 0.25 | YY a*a | M |  |  |
| F4 x M2 | 0.125 | XX a*a | F | 0.625 F |  |
|  | 0.25 | XY a*a | M | 0.375 M |  |
|  | 0.125 | YY a*a | M |  |  |
|  | 0.125 | XX a*a* | F |  |  |
|  | 0.25 | XY a*a* | F |  |  |
|  | 0.125 | YY a*a* | F |  |  |
| F4 x M3 | 0.25 | XY a*a | M | 0.5 F |  |
|  | 0.25 | YY a*a | M | 0.5 M |  |
|  | 0.25 | XY a*a* | F |  |  |
|  | 0.25 | YY a*a* | F |  |  |
| F5 x M1 | 0.25 | XY a*a | M | 1 M |  |
|  | 0.25 | YY a*a | M |  |  |
| F5 x M2 | 0.25 | XY a*a | M | 0.5 F |  |
|  | 0.25 | YY a*a | M | 0.5 M |  |
|  | 0.25 | XY a*a* | F |  |  |
|  | 0.25 | YY a*a* | F |  |  |
| F5 x M3 | 0.5 | YY a*a | M | 0.5 F | NOTE: recurrent genotype pair in rainbow darters |
|  | 0.5 | YY a*a* | F | 0.5 M |  |

**Extended Data Table 3.** SIFT predictions for mutations in sex-associated mitonuclear genes.

| Gene | Chromosome | Substitution | Derived | SIFT score | Prediction |
| --- | --- | --- | --- | --- | --- |
| *ndufa7* | 9 | H18R | *E. spectabile* | >0.05 | Tolerated |
| *ndufa11* | 9 | V44M | *E. caeruleum* | >0.05 | Tolerated |
| *ndufb5* | 9 | N24S | *E. spectabile* | >0.05 | Tolerated |
|  |  | P35A | *E. caeruleum* | >0.05 | Tolerated |
| *ndufs7* | 9 | P45S | *E. spectabile* | 0.04 | Not tolerated |
| *ndufa12* | 23 | Q116R | *E. spectabile* | 0.02 | Not tolerated |
|  |  | Q124K | *E. caeruleum* | >0.05 | Tolerated |
| *mt-nd2* | MT | A8T | *E. caeruleum* | >0.05 | Tolerated |
|  |  | I19V | *E. spectabile* | >0.05 | Tolerated |
|  |  | L70I | *E. spectabile* | 0.02 | Not tolerated |
|  |  | T90S | *E. spectabile* | >0.05 | Tolerated |
|  |  | I95T | *E. caeruleum* | >0.05 | Tolerated |
|  |  | L97V | *E. spectabile* | 0.04 | Not tolerated |
|  |  | I98V | *E. spectabile* | >0.05 | Tolerated |
|  |  | T99I | *E. caeruleum* | >0.05 | Tolerated |
|  |  | A148T | *E. spectabile* | >0.05 | Tolerated |
|  |  | S150T | *E. caeruleum* | >0.05 | Tolerated |
|  |  | L212F | *E. caeruleum* | >0.05 | Tolerated |
|  |  | N221S | *E. spectabile* | >0.05 | Tolerated |
|  |  | T224A | *E. caeruleum* | >0.05 | Tolerated |
|  |  | A236T | *E. caeruleum* | >0.05 | Tolerated |
|  |  | A270T | *E. caeruleum* | >0.05 | Tolerated |
|  |  | L308I | *E. caeruleum* | >0.05 | Tolerated |
|  |  | T309A | *E. caeruleum* | >0.05 | Tolerated |
|  |  | T328A | *E. spectabile* | >0.05 | Tolerated |
|  |  | T339A | *E. caeruleum* | >0.05 | Tolerated |
| *mt-nd3* | MT | I8V | *E. spectabile* | >0.05 | Tolerated |
|  |  | V9L | *E. spectabile* | >0.05 | Tolerated |
|  |  | I16V | *E. spectabile* | >0.05 | Tolerated |
|  |  | A83V | *E. spectabile* | >0.05 | Tolerated |
|  |  | I104V | *E. spectabile* | >0.05 | Tolerated |
| *mt-nd5* | MT | S5A | *E. caeruleum* | >0.05 | Tolerated |
|  |  | I14V | *E. spectabile* | 0.05 | Not tolerated |
|  |  | T17S | *E. spectabile* | >0.05 | Tolerated |
|  |  | A20T | *E. caeruleum* | >0.05 | Tolerated |
|  |  | F21L | *E. caeruleum* | >0.05 | Tolerated |
|  |  | G33K | *E. spectabile* | >0.05 | Tolerated |
|  |  | H34P | *E. caeruleum* | >0.05 | Tolerated |
|  |  | L38I | *E. caeruleum* | >0.05 | Tolerated |
|  |  | E62D | *E. caeruleum* | 0.02 | Not tolerated |
|  |  | T66A | *E. caeruleum* | >0.05 | Tolerated |
|  |  | I68V | *E. caeruleum* | >0.05 | Tolerated |
|  |  | M74I | *E. caeruleum* | 0.03 | Not tolerated |
|  |  | H89F | *E. caeruleum* | >0.05 | Tolerated |
|  |  | I97V | *E. spectabile* | >0.05 | Tolerated |
|  |  | A115T | *E. spectabile* | >0.05 | Tolerated |
|  |  | F118Y | *E. caeruleum* | >0.05 | Tolerated |
|  |  | V128T | *E. spectabile* | >0.05 | Tolerated |
|  |  | I135V | *E. caeruleum* | >0.05 | Tolerated |
|  |  | A167T | *E. spectabile* | >0.05 | Tolerated |
|  |  | M194V | *E. caeruleum* | 0.02 | Not tolerated |
|  |  | V206L | *E. spectabile* | >0.05 | Tolerated |
|  |  | D212G | *E. caeruleum* | >0.05 | Tolerated |
|  |  | F213Y | *E. caeruleum* | >0.05 | Tolerated |
|  |  | P277S | *E. caeruleum* | >0.05 | Tolerated |
|  |  | C378S | *E. caeruleum* | 0.01 | Not tolerated |
|  |  | H440Y | *E. spectabile* | >0.05 | Tolerated |
|  |  | N444S | *E. spectabile* | >0.05 | Tolerated |
|  |  | V469I | *E. caeruleum* | >0.05 | Tolerated |
|  |  | L472F | *E. spectabile* | >0.05 | Tolerated |
|  |  | T475S | *E. spectabile* | >0.05 | Tolerated |
|  |  | P489T | *E. spectabile* | 0.02 | Not tolerated |
|  |  | L494I | *E. spectabile* | 0.01 | Not tolerated |
|  |  | N515S | *E. caeruleum* | >0.05 | Tolerated |
|  |  | T538L | *E. caeruleum* | 0.04 | Not tolerated |
|  |  | L543V | *E. spectabile* | >0.05 | Tolerated |
|  |  | M573I | *E. caeruleum* | >0.05 | Tolerated |
|  |  | T578A | *E. caeruleum* | >0.05 | Tolerated |
|  |  | R589Q | *E. spectabile* | >0.05 | Tolerated |
|  |  | I592M | *E. spectabile* | >0.05 | Tolerated |
|  |  | A597S | *E. spectabile* | >0.05 | Tolerated |
|  |  | L609V | *E. spectabile* | >0.05 | Tolerated |
|  |  | F610L | *E. caeruleum* | >0.05 | Tolerated |
|  |  | T611A | *E. spectabile* | >0.05 | Tolerated |

*In silico* functional predictions from SIFT for variants in nuclear genes that encode complex I subunit proteins present on *E. spectabile* and *E. caeruleum* sex chromosomes (9 and 23, respectively) and variants in mitochondrial genes that are associated with complex I and are under selection in these species<sup>42</sup>. Variants are indicated as being derived in *E. spectabile* or *E. caeruleum*. A SIFT score of >0.05 is expected to be tolerated, and 0.05 or less is expected to be not tolerated and potentially have deleterious effects on protein function.

