## Supplementary Methods for "Sex chromosome turnover and mitonuclear conflict drive reproductive isolation"

Etheostoma caeruleum *reference genome annotation*

We assessed the completeness of the rainbow darter assembly with BUSCO (Benchmarking Universal Single-Copy Orthologs; v5.7.1), specifying the Actinopterygii_odb10 database that contains a curated set of conserved single-copy orthologs present in ray-finned fishes^1^. The presence, absence, or fragmentation of these genes gives us a quantitative estimate of how complete the genome is in terms of expected gene content. In addition, we ran QUAST v5.2.0^2^ to generate standard assembly statistics (e.g., number of contigs, N50).

We masked repetitive material in the genome (such as regions encoding transposable elements) prior to gene annotation. We ran RepeatModeler v2.0.5^3^ to detect and define repetitive regions in the genome and then ran RepeatMasker v4.1.4^4^ to mask identified repeat regions. To define coordinates for genes, exons, and other coding and non-coding sequences in the assembly, we ran BRAKER3^5^ with AUGUSTUS gene prediction^6^. BRAKER3 was selected as it makes predictions based on both RNA and protein input evidence. We first downloaded the *de novo* transcriptome sequence data of a female yellow perch (*Perca flavescens*, accession number SRR3498542), a close relative of the darter clade, and the transcriptome of *Etheostoma spectabile* (accession number SRR9855987) using the fasterq-dump command in SRA-Toolkit (v3.1.1) from NCBI (SRA Toolkit Development Team 2020). These raw paired-end RNA-seq reads were then aligned to our *Etheostoma caeruleum* genome assembly using STAR v2.7.10b^7^. Resulting alignment files were used as transcriptomic evidence for BRAKER3. To enhance prediction accuracy, we provided a custom protein database containing annotated protein sequences across Actinopterygii. These protein sequences served as external evidence and were used by the ProtHint pipeline within BRAKER3 to generate intron and exon hints, guiding AUGUSTUS in the identification of gene structures. Finally, we provided the resulting annotation file from BRAKER3 to eggNOG-mapper v2 for functional identification of sequences identified as genes^8,9^.

*Quantifying introgression on the sex chromosomes and autosomes*

We used topology weighting and calculated ABBA-BABA statistics across the genome to ask whether sex-linked regions are more resistant to introgression between *E. spectabile* and *E. caeruleum* compared to autosomes. Such a pattern would further support that sex chromosomes play a role in reproductive isolation. We first inferred tree topologies across the genome to identify regions harboring discordant topologies indicative of introgression. For this analysis, we included allopatric *E. spectabile* (Sangamon River, IL, n=24, allopatric *E. caeruleum* (Kalamazoo River, MI, n=14), sympatric *E. spectabile* and *E. caeruleum* (Vermillion River, IL, n=24 each), and an outgroup (fantail darters, *E. flabellare*, Sangamon River, IL, n=11) (see Extended Data Table 1 for collection location coordinates). Phyml was used to build trees in 50 SNP windows across each chromosome and input to TWISST (Topology Weighting by Iterative Sampling of Sub-trees)^15^. Given that we included five taxa in the analyses, a total of 15 unique topologies were possible. TWISST was used to assign weightings to each of the 15 topologies at each 50 SNP window across the genome, with topology 11 representing the species tree (Extended Data Fig. 6). To compare the rankings of topologies within a given region of interest (i.e., significantly sex-associated regions on sex chromosomes) we normalized the rankings for each 50 SNP window to proportions ranging from 0 to 1. Thus, topologies with higher weightings for a given window are represented with rankings closer to 1.

We next conducted ABBA-BABA derived tests for introgression from sympatric *E. spectabile* into sympatric *E. caeruleum* and vice versa. For this analysis, we used computationally phased, biallelic SNPs to quantify whether allele frequencies follow those expected between three lineages (e.g., sister lineages P1 and P2, and a third closely related lineage, P3) under expectations for incomplete lineage sorting (ILS). Observing a greater proportion of shared derived alleles between P2 and P3 (ABBA) or between P1 and P3 (BABA) than what would be expected by chance (i.e., ILS) indicates introgression. The fantail darter *E. flabellare* served as an outgroup to allow for polarizing ancestral versus derived alleles. Using the script ABBABABAwindows.py (https://github.com/simonhmartin/genomics_general/), we calculated *fdM* in 20 kb non-overlapping windows across the genome, requiring a minimum of 10 SNPs per window. *fdM* is bounded by -1 and 1 with positive values indicative of introgression between P1 and P3 and negative values indicative of introgression between P2 and P3. The *fdM* statistic is also not as sensitive to reduced variation within populations as other ABBA-BABA tests^92^. We used Wilcoxon rank sum test to compare the mean *fdM* for sex chromosomes versus autosomes.

Lastly, we investigated whether genomic regions with stronger sex-association within species, indicated by lower p-values from the GEMMA GWAS analyses, also exhibited higher weighting for the species tree topology (topology 11, Extended Data Fig. 6). A positive correlation would suggest that regions showing strong divergence between the sexes within species are also resistant to introgression between species. To test this, we performed a Pearson’s correlation between the species tree topology weightings and GWAS p-values for chromosomes 9 and 23. We identified genomic windows with both the most significant GWAS p-values and the highest species tree topology weights, aligned to the *E. spectabile* reference genome.

**Supplementary Note**

*Identifying candidate sex determining genes in* Etheostoma radiosum

We conducted whole genome resequencing on three closely related species within the orangebelly darter (*E. radiosum*) complex, including *E. radiosum radiosum*, *E. radiosum cyanorum*, and *E. radiosum paludosum*^16^ (see Extended Data Table 1). Initial separate analyses on each of these species showed consistent patterns pointing to chromosome 22 as the sex chromosome. Accordingly, to provide increased power to detect sex linked loci, we pooled samples for all *E. radiosum* spp. for subsequent analyses. We identified two highly significant GWAS regions for sex on chromosome 22 in the *E. radiosum* complex, with a first larger peak spanning 9.8 - 11.8 Mb and a second narrow peak at 14.4 Mb. We identified 86 genes within 50 kb of these significantly sex-associated regions using the *E. spectabile* genome annotation. GO enrichment analysis revealed these genes were enriched for ontologies including negative regulation of luteinizing hormone secretion, negative regulation of gonadotropin secretion, pituitary gland development, adenohypophysis development, regulation of mitochondrial DNA replication, regulation of mitochondrial DNA metabolic process, and regulation of mitochondrial gene expression (Table S3). Within the sex-associated stratum on chromosome 22 in *E. radiosum*, we identified *shh* (sonic hedgehog), a key developmental regulator implicated in the formation of the pituitary gland^17^, and *oprk1*, which encodes the kappa-opioid receptor known to suppress GnRH/LH secretion via hypothalamic pathways^18^. The presence of these genes within a putative sex-determining region suggests a potential mechanistic link between sex chromosome divergence and hormonal regulation of reproductive function. Two additional genes within the sex-associated region on chromosome 22, *ptprn2* and *adcyap1*, are also linked to neuroendocrine signaling. *Ptprn2* encodes a tyrosine phosphatase involved in neurosecretory pathways^19^, while *adcyap1* encodes a neuropeptide that regulates pituitary hormone release^20^, including growth hormone. In medaka, *adcyap1* shows extreme male-biased expression in the preoptic area of the brain, with these differences driven by sex-specific estrogen levels and hormonal environments^21^ , paralleling potential mechanisms of sex-biased pituitary regulation in darters. This points to a potential role for pituitary axis modulation in sex differentiation in the *E. radiosum* species complex.

Notably, the GWAS peak at 14.4 Mb lies within ~500 kb of *ndufb4*, a nuclear-encoded subunit of mitochondrial Complex I. NDUFB4 interacts with NDUFS7, NDUFB5, and MT-ND5 in complex I, and these genes have been shown to harbor derived and positively selected variants in *E. spectabile* and *E. caeruleum*^22^ (Extended Data Table 3). These results add to a pattern across darter species in which independently evolved sex chromosomes consistently harbor mitonuclear genes. This raises the possibility that mitonuclear conflict, particularly in hybridizing lineages, may favor the recruitment of alternative sex chromosomes that restore cytonuclear compatibility.

**Supplementary Table Legends**

**Supplementary Table 1. EggNOG-mapper functional annotation of 10X rainbow darter reference genome.** Functional annotation of the 10X *E. caeruleum* reference genome using EggNOG-mapper. This tool uses orthology predictions based on the eggNOG database to assign functional information to genes predicted in gene annotation software (such as BRAKER3, used for this genome).

**Supplementary Table 2. Summary of statistical tests for sex chromosome vs. autosome comparisons.** Wilcoxon rank-sum test results are summarized, including W statistic, sample sizes, mean differences, Cliff’s δ, and 95% confidence intervals for analyses of male – female difference in heterozygosity and SNP density on sex chromosome vs. autosomes,

**Supplementary Table 3. Candidate genes near sex-associated loci and divergence outliers in orangethroat and rainbow darters.** Gene names and coordinates for all genes within 50 Kb of significantly sex-associated SNPs in *E. spectabile* (Espec), *E. caeruleum* (Ecaer), and *E. radiosum* (Erad), and intersex Fst gene outliers (top 5% Fst out of all genes in the annotation) for allopatric and sympatric populations of *E. spectabile* and *E. caeruleum.*

**Supplementary Table 4. Enriched gene ontologies for outliers in sex association.** Results of the Gene Ontology (GO) enrichment analyses on sets of highly sex-associated genes (GWAS p-value <5e-8) within *E. spectabile* (Espec), *E. caeruleum* (Ecaer), and *E. radiosum* (Erad). Enrichment analyses were conducted with both zebrafish and human as the reference.

**Supplementary Table 5. Introgression statistics for loci across the genome.** Results of ABBA-BABA analyses in 20 kb windows across the genome. *SymEc2SymEs_introgression_stats*: Test for introgression from sympatric *E. caeruleum* into sympatric *E. spectabile. SymEs2SymEc_introgression_stats*: Test for introgression from sympatric *E. spectabile* into sympatric *E. caeruleum.*

**Supplementary Table 6. Genome-wide distribution of SNPs associated with mitochondrial background.** SNPs with mitonuclear association across the genome. Chromosomes 9, 23, and 18 have the most SNPs significantly associated with mitochondrial background. *ChiSqMitonucAssoc_deltaMAF_bx*: Chi square tests and FDR-adjusted p-value for deviations from the expected delta MAF for each SNP. *FETsigSNPs_perLG_mitonuc_bx:* Fisher’s Exact Tests asking whether there is an enrichment of significant SNPs above that expected by chance on each chromosome**.** *MitoNuc_Assoc_Genes_GO_danio*: Results of GO enrichment analysis on genes with a significant mito-nuclear association in backcross hybrids, with the zebrafish genome as the reference. *MitoNuc_Assoc_Genes_GO_human:* Results of GO enrichment analysis on genes with a significant mito-nuclear association in backcross hybrids, with the human genome as the reference.
